# Engineered SMCHD1 and D4Z4 mutations reveal roles of D4Z4 heterochromatin disruption and feedforward DUX4 network activation in FSHD

**DOI:** 10.1101/2022.10.14.512332

**Authors:** Xiangduo Kong, Nam Viet Nguyen, Yumeng Li, Jasmine Shaaban Sakr, Kate Williams, Sheila Sharifi, Jonathan Chau, Altay Bayrakci, Seiya Mizuno, Satoru Takahashi, Tohru Kiyono, Rabi Tawil, Ali Mortazavi, Kyoko Yokomori

**Affiliations:** Department of Biological Chemistry, School of Medicine, University of California, Irvine, CA; Department of Development and Cell Biology, School of Biological Sciences, University of California, Irvine, CA; Laboratory Animal Resource Center in Transborder Medical Research Center, University of Tsukuba, Tsukuba, Ibaraki, Japan; Department of Anatomy and Embryology, Faculty of Medicine, University of Tsukuba; Division of Carcinogenesis and Cancer Prevention, National Cancer Center Research Institute, Tsukiji, Chuo-ku, Tokyo, Japan; Neuromuscular Disease Unit, Department of Neurology, University of Rochester Medical Center, Rochester, New York, USA

## Abstract

Facioscapulohumeral dystrophy (FSHD) is commonly associated with contraction of D4Z4 repeats on chromosome 4q (FSHD1). Mutations in the *SMCHD1* gene are linked to both minor cases with no prominent repeat loss (FSHD2) and severe cases of FSHD1. Abnormal upregulation of the transcription factor DUX4, encoded in the *D4Z4* repeat, is believed to play a central role in FSHD. However, defining the disease mechanism has been hampered by the heterogeneity of patient-derived cells, difficulty to detect DUX4 in patient myocytes, and limited animal models because D4Z4 repeats are primate-specific. To overcome these limitations, we engineered isogenic human skeletal myoblast lines with D4Z4 and/or *SMCHD1* mutations. We found a highly synergistic effect of double mutations on triggering two key disease processes, D4Z4 heterochromatin disruption and cross-stimulation of DUX4 targets, such as histone H3.X/Y and LEUTX transcription factor. Thus, engineered human myocyte models provide unique insights into the molecular mechanisms underpinning FSHD.

**Teaser:** FSHD mutations cause D4Z4 heterochromatin disruption and feedforward DUX4 network activation.

## Introduction

Facioscapulohumeral dystrophy (FSHD) is one of the most common muscular dystrophies with a prevalence of ~1 in 8,333. FSHD causes progressive wasting of facial, shoulder, and upper arm as well as lower leg musculature (1, 2). The majority of FSHD cases (>95%) are caused by monoallelic contraction of 3.3 kb D4Z4 macrosatellite repeat sequences located at the subtelomeric region of chromosome 4q (4qter D4Z4) (FSHD1 (MIM 158900)) (3, 4). FSHD1 is associated with 1~10 copies of D4Z4 repeats in the contracted allele in contrast to 11~150 copies in the intact allele. Lower copy numbers (1~3 copies) are tied to earlier onset and more severe phenotypes compared to higher copy numbers (8~10 copies). However, clinical manifestations are variable, suggesting a potential contribution of additional modifier gene(s) (2, 5). FSHD2, which is the rare form of FSHD (<5% of cases), is mainly linked to mutations of *SMCHD1* (MIM 158901) (2, 6). Although FSHD2 was thought previously to involve no D4Z4 repeat contraction, accumulating evidence indicates its association with relatively short D4Z4 repeats (8~20 repeats) (2). Moreover, mutations of the *SMCHD1* gene have been found in severe cases of FSHD1, suggesting that it may act as a modifier gene to increase the disease severity (7, 8).

D4Z4 contains an open reading frame for the double-homeobox transcription factor *DUX4* gene (9–11). *DUX4* gene is normally expressed early in embryogenesis and participates in zygotic genome activation (12, 13). Expression of short and long isoforms have been detected but only the full-length *DUX4* transcript (*DUX4fl*) that includes a transactivation domain can activate target genes, and its abnormal upregulation is associated with FSHD (11, 14, 15). The *DUX4* gene, embedded in the D4Z4 repeat, lacks a poly-adenylation (poly(A)) signal sequence, and only those individuals with “permissive” 4qA haplotypes carrying the canonical poly(A) signal downstream of the last D4Z4 repeat develop FSHD, strongly suggesting that the disease is tightly linked to functional *DUX4* mRNA production from the last D4Z4 copy (14). Indeed, upregulation of DUX4 target genes is readily detectable in patient cells, supporting the significance of DUX4-mediated gene activation in FSHD. Curiously, however, *DUX4fl* RNA and protein expressions are extremely infrequent and occur at low levels in patient myocytes, albeit higher than in control cells (11, 14–16). Consequently, expression of DUX4 target genes, rather than *DUX4* itself, is used as markers for the FSHD phenotype (5, 10, 17–20). However, dynamics and regulation of the DUX4 target gene network activation is not well understood.

Hypomethylation of D4Z4 DNA is a signature change in FSHD patient cells (21–23). We also found that D4Z4 repeats contain heterochromatic regions marked by histone H3 lysine 9 trimethylation (H3K9me3), heterochromatin binding protein HP1γ and the higher-order chromatin organizer cohesin (24). This heterochromatin structure is compromised in both FSHD1 and FSHD2 (24). Indeed, reduction of H3K9me3 at D4Z4 by inhibition or depletion of SUV39H1 causes *DUX4fl* expression (24, 25). Thus, perturbation of heterochromatinization of D4Z4 appears to be directly linked to FSHD pathogenesis (15, 26). SMCHD1 binds to D4Z4, and its haploinsufficiency results in derepression of *DUX4fl* expression, indicating a direct role of SMCHD1 in *DUX4fl* regulation (6). We found that SMCHD1 binding to D4Z4 is H3K9me3-dependent (25). This raised the possibility that even in FSHD1 with no mutation in *SMCHD1*, SMCHD1 binding to D4Z4 may be compromised (due to the loss of H3K9me3), which may contribute to *DUX4* upregulation (25). In severe cases of FSHD1, this effect may be further exacerbated by the actual mutations in *SMCHD1* itself (7, 8). SMCHD1 has been implicated in regulation of DNA methylation at certain CpG islands and at the inactive X chromosome (27–29), and thus, it is assumed to also regulate DNA methylation at D4Z4. However, SMCHD1 represses gene expression in both DNA methylation-dependent and independent ways (30), and whether SMCHD1 modulates DUX4fl expression through DNA methylation has not been determined.

Overexpression of recombinant DUX4 in *in vitro* myoblasts and in *in vivo* model organisms is highly toxic (31, 32), and there is an ongoing effort to characterize the mechanism of DUX4-induced cell toxicity (33–36). However, the frequency of DUX4 expression in patient cells is often less than 1%, which is substantially lower than that in the recombinant DUX4-inducible systems. In fact, recent single cell/nucleus-sequencing and in situ RNA detection analyses revealed no significant evidence for cell death in FSHD patient myocytes that endogenously express DUX4 (37, 38). Moreover, the subcellular localization of the recombinant *DUX4fl* mRNA differs from that of the endogenous *DUX4fl* mRNA (37, 39). Collectively, these observations raise the question whether DUX4-induced acute cytotoxicity is a physiologically relevant mechanism of disease pathogenesis. To circumvent these issues, efforts are being made to express DUX4 at low level in an inducible fashion in mice (40, 41). D4Z4 repeats (and *DUX4* within), however, are primate-specific. Likewise, DUXA and LEUTX, two main transcription factors activated by DUX4, are absent in mice (42). Consequently, the genes and molecular network activated by human *DUX4* introduced in non-primate model organisms are different from those in patient muscle. Thus, a physiologically relevant model for FSHD is still lacking.

Crucially, whether genetic changes (D4Z4 contraction and/or mutations in *SMCHD1*) are sufficient to recapitulate patient phenotypes remains undetermined. To experimentally address this and to further investigate the disease mechanism, here, we report the development of CRISPR-engineered isogenic mutant myoblast cell lines carrying either D4Z4 mutations, SMCHD1 mutation or both. Analyses of these cell lines provided evidence for a feedback loop between SMCHD1 and H3K9me3 at D4Z4 and demonstrated a synergistic effect of double mutations on DUX4 target gene induction. Additional heterochromatin disruption enhanced the effect of individual mutations. Furthermore, our results uncovered differentiation-insensitive early and -sensitive late DUX4 target genes and their positive cross-regulation. Taken together, our analyses of isogenic FSHD mutant myocytes reveal heterochromatin disruption and feedforward gene expression network downstream of DUX4 as two key processes underlying FSHD pathogenesis.

## Results

### Elimination of SMCHD1 has a minor effect on *DUX4* expression

FSHD occurs only in individuals with “permissive” 4qA haplotypes, in which the presence of a poly(A) signal downstream of the last D4Z4 copy would allow the expression of functional DUX4fl mRNA (14). To compare the effects of haplotypes, SMCHD1 shRNA depletion was initially performed in healthy control cell lines with non-permissive and permissive haplotypes (Supplemental Fig. S1). Interestingly, in both cell types, this caused upregulation of DUX4 target genes at the myoblast stage in a statistically significant and comparable fashion compared to control shRNA-treated cells, suggesting that a low level functional *DUX4* mRNA can be expressed in non-permissive cells (Supplemental Fig. S1A). Notably, DUX4 target gene expression was further stimulated upon myotube differentiation in permissive haplotype cells. In contrast, despite comparable SMCHD1 depletion efficiency, no further stimulation was observed in non-permissive haplotype myotubes (Supplemental Fig. S1B). Thus, the permissive haplotype is critical for differentiation-induced stimulation of DUX4 and target gene expression.

Even in cells with a permissive haplotype, DUX4 target gene expression in SMCHD1-depleted myotubes is much lower than that in FSHD2 myotubes (Supplemental Fig. S1C). Thus, transient SMCHD1 depletion is not sufficient for a robust FSHD phenotype. To generate stable mutant cell lines, gRNAs specific for *SMCHD1* were designed for CRISPR knockout (KO) (Supplemental Fig. S2A). Despite screening 300 clones, we failed to obtain heterozygous mutant cells due to high efficiency of CRISPR mutation. Thus, unlike FSHD2 cells, in which *SMCHD1* mutation is heterozygous, our *SMCHD1* (SM) mutant cells are *SMCHD1* null (Fig. 1A). As in HCT116 colorectal cancer cells (43), *SMCHD1* KO in adult myoblasts is not lethal. *SMCHD1* mutation in permissive haplotype cells upregulated DUX4 target genes in a statistically significant fashion compared to the parental control cells (Fig. 1D). However, even after a complete loss of SMCHD1, the amount of DUX4 target gene transcripts was much lower than that in FSHD2 patient cells (which harbor a heterozygous *SMCHD1* mutation). Thus, by itself, a somatic KO mutation of *SMCHD1* is not sufficient to recapitulate FSHD2 (Fig. 1E).

**Figure 1.**
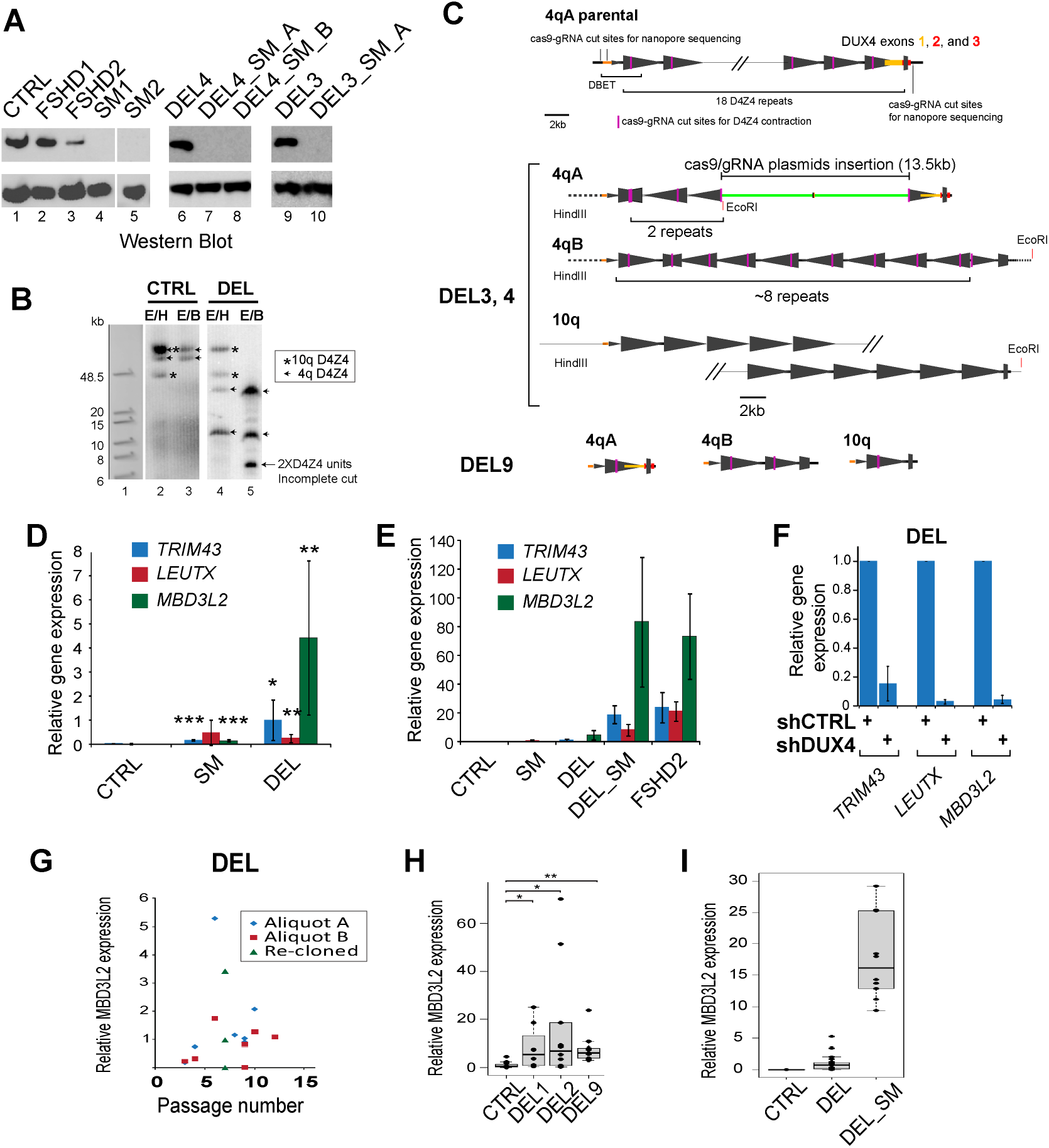
Generation of SMCHD1 and/or D4Z4 mutant cells from health permissive skeletal myoblast. **A.** Western blot analysis the SMCHD1 protein expression in the cell lines used in the study. Lysates of immortalized control and FSHD1 and FSHD2 patient myoblasts, SMCHD1 mutants (SM), D4Z4 deletion mutants (DEL) and double mutants (DEL_SM) were subjected to western blot analysis using antibody specific for SMCHD1. β-Tubulin serves as a loading control. **B.** Determination of the 4q and 10q D4Z4 repeat number. Examples of control and DEL mutant clones are shown. Genomic DNA was digested with EcoRI/HindIII (E/H) or EcoRI/BlnI (E/B) and subjected to PFGE. They were then blot-hybridized with the 4q/10q specific “1-kb” D4Z4 probe. E/H digestion leaves intact two 4q and two 10q D4Z4 arrays, while BlnI in an E/B only cleaves 10qD4Z4 repeat units. Size markers (in kb) are shown on the left. Arrowheads and stars indicate 4q and 10q D4Z4, respectively. The arrow indicates a band around 6.6kb, which should be 2 D4Z4 repeat units caused by incomplete digestion. The two 4q D4Z4 repeat arrays are contracted, while the 10q D4Z4 bands size show no change. **C.** D4Z4 gRNA targeting resulted in repeat contraction and recombination, leaving the last repeat with the *DUX4* gene intact at the 4qA allele in DEL mutant cells. Top: schematic diagram of D4Z4 array in 4qA allele of parental cell with gRNA target sites for D4Z4 deletion (purple bars) as well as crRNA target sites designed for nanopore sequencing were shown at the top panel. D4Z4 cluster in 4qA, 4qB and 10q alleles of DEL3 were shown below. 10q D4Z4 sequences were confirmed by SNP analysis. The large triangle represented a 3.3kb D4Z4 unit and its orientation. The small and partial triangle represented partial D4Z4 units and their orientation. The endonucleases (EcoRI/HindIII) cut sites, which generated the fragments detected in PFGE, are indicated. **D.** Control, SMCHD1 mutant SM1 and D4Z4 contraction mutant DEL3 myoblasts were differentiated and analyzed for DUX4 target (*TRIM43, LEUTX* and *MBD3L2*) RNA expression levels. Data are expressed as relative expression (mean with standard deviation). The gene expression over *GAPDH* was normalized to the *TRIM43* value of DEL, which was set to be 1. * P < 0.05, **P < 0.01 and ***P < 0.001 vs. control. **E.** Corresponding data from double mutant (DEL_SM) and FSHD2 were added to (D) for comparison. **F.** DUX4 depletion by lentiviral shRNA abolished activation of DUX4 target genes (*TRIM43, LEUTX* and *MBD3L2*) compared to control shRNA (shCTRL). Cells were harvested at 4 days of myotube differentiation. DUX4 target expression levels were determined by RT-qPCR. Y-axis is relative expression (mean with standard deviation) with the expression in shCTRL-transduced samples as one. **P < 0.01 and ***P < 0.001. **G.** Stochasticity of target gene expression in DEL3. Two aliquots of the same clone at different passage numbers and replicates of a re-cloned cell line were differentiated for 4 days and analyzed for *MBD3L2* expression by RT-qPCR. **H.** Variegation is also observed in multiple DEL mutant clones, including DEL9 as in (**G)**. **I.** Comparison of *MBD3L2* expression level in early myotube of Control (n=11), single mutant DEL (n=23), and double mutant DEL_SM (n=10). The dots on each boxplot represent the individual data in each repeat.

### D4Z4 contraction induces variegated DUX4 target gene expression, which is enhanced by concurrent loss of SMCHD1

A set of gRNAs was designed at the single-nucleotide polymorphism (SNP) regions to enhance mutations of 4q over 10q D4Z4 repeats (DEL mutants) (Fig. 1A-C and Supplemental Fig. S2B). Parental control cells have 24 and 18 copies of D4Z4 on chromosome 4q and 24 and 12 copies on chromosome 10q based on PFGE (Fig. 1B). We found that CRISPR-CAS9-mediated D4Z4 disruption was sufficient to upregulate DUX4 target genes (e.g., TRIM43, LEUTX and MBD3L2) in early myotubes, which was therefore used as a screening phenotype (Fig. 1D and Supplemental Fig. S2B). We generated DEL mutant clones using DNA plasmid- and protein/RNA-based CRISPR-Cas9 systems by screening ~300 and 180 clones, respectively (see Methods and Supplemental Figure S2B). We observed a tendency of inverse correlation between high DUX4 target gene expression and efficient myotube differentiation (Supplemental Fig. S2C). For our experiments, we selected clones with a relatively high DUX4 target gene expression as well as efficient proliferation and differentiation capabilities (Supplemental Fig. S2C). We found that D4Z4 mutant (DEL) clones showed higher DUX4 target gene expression than SM clones, but still much lower than FSHD patient cells (DEL3 as an example in Fig. 1D and E). Importantly, target gene upregulation is DUX4-dependent as DUX4 shRNA depletion abolished the expression of target genes (Fig. 1F). We found that target gene expression is highly variable even in the same clone with different passage numbers and/or in different experiments (Fig. 1G).

Although PFGE analyses indicated repeat contraction (Fig. 1B), we analyzed the regions and resulting transcripts using genomic and RNA nanopore long-read sequencing, respectively. For DEL mutants generated by the plasmid-based CRISPR-CAS9 treatment, we found that gRNA-mediated cutting of D4Z4 repeats in the 4qA allele resulted not only in deletion of repeats upstream of the last copy, but also in insertion of inverted 2.5 copies of D4Z4 repeats as well as 2 copies of a gRNA plasmid sequence separated by a small fragment of a CAS9 plasmid sequence (Fig. 1C; Supplemental Fig. S2D). These inverted repeats could give rise to the ~2 repeat signal in PFGE when in fact only the last copy was left downstream of this insertion, thus creating the 4qA allele with one D4Z4 repeat (Fig. 1B). For 4qB, we found the repeat shortening and inversion, the length consistent with the PFGE band, which would not yield any significant DUX4 gene expression due to the absence of canonical poly(A) signal sequence (14) (Fig. 1C). No rearrangements were detected in 10q D4Z4 alleles, consistent with the expectation with 4q-tailored gRNAs (Fig. 1C). The insertion of plasmid sequences resulted in expression of EGFP from the PKG promoter as well as small spliced RNA fragments containing *DUX4* exons 2 and 3 (corresponding to the 3’-UTR of *DUX4* mRNA) (termed a chimeric DUX4 3’-UTR) (Supplemental Fig. S2D). We confirmed that overexpression of the chimeric DUX4 3’UTR fragment has no effect on DUX4 target gene expression (Supplemental Fig. S2E). In contrast, protein/RNA-based CRISPR-Cas9 mutagenesis resulted in repeat contraction down to one copy of D4Z4 at 4qA and 2 copies at 4qB, and also reduced the copy number of 10qD4Z4 alleles to one copy as confirmed by nanopore sequencing (Fig. 1C, DEL9). DEL9 cells exhibit comparable upregulation of DUX4 target genes as the plasmid-based DEL mutants (Fig. 1H, *MBD3L2* as an example). Consistent with the one intact *DUX4* gene in the 4qA allele, RNA nanopore sequencing of both types of DEL mutant clones confirmed an intact wild type *DUX4fl* transcript, indicating that the last D4Z4 copy at 4qA retained the ability to express *DUX4fl* (Supplemental Fig. S2D). Thus, we used both of these mutant clones for further analyses.

The above results demonstrate that D4Z4 mutations cause upregulation of *DUX4* and target genes, but their expression is highly variegated and tends to be much lower than in FSHD patient cells (Fig. 1E). In FSHD1 patients, the lower D4Z4 repeat numbers associate with more severe clinical phenotypes (2, 44). Our results suggest, however, that even when only one repeat copy left, additional mechanism(s) are required to recapitulate the full FSHD-associated gene expression program. As *SMCHD1* mutations are associated with severe cases of FSHD1 (combined with D4Z4 contraction) (7, 8), we generated double mutant cells (DEL_SM), by introducing *SMCHD1* mutation in DEL clones (Supplemental Fig. S2B). In comparison to single DEL mutants, DEL_SM mutants consistently upregulated DUX4 target genes at a higher level (Fig. 1E and I). These results demonstrate that *SMCHD1* acts as a modifier gene whose loss acts synergistically with D4Z4 contraction to enhance and stabilize the DUX4 target gene expression.

### Synergistic effect of double mutations recapitulates patient cell phenotype

To determine the effects of these engineered mutations, we analyzed the genome-wide gene expression changes during myoblast differentiation by RNA-seq of 3-5 clones of each mutant type (SM, DEL and DEL_SM) in comparison to the isogenic control as well as FSHD1 and FSHD2 patient cells. Although we chose clones with relatively efficient differentiation, we observed prominent delays in differentiation of DEL and DEL_SM clones even in those with comparable doubling time (Supplemental Fig. S3A). A mild delay was also observed in FSHD patient cells whereas the delay was minimal in *SMCHD1* mutant (SM) cells. Time course principal component analyses (PCA) of gene expression during differentiation also confirmed delays in DEL and DEL_SM mutant cells compared to the isogenic control and SM cells (Supplemental Fig. S3B). To compensate for the differentiation delay, we compared DEL and DEL_SM samples at days 4~5 and 13~14 of with control and SM samples on days 3 and 12 of differentiation as “early” and “late” myotubes, respectively, in the current study.

Using edgeR, we identified up- and downregulated differential expressed genes (DEGs) (*P*-values <0.01 and Log_2_ fold change > 1.5) in each mutant type (Supplemental Table S1). Interestingly, at both myoblast and early myotube stages, the number of genes that are upregulated in the DEL_SM mutant cells is significantly more (673 and 1228, respectively) than the sum of de-repressed genes in the DEL- or SM-only mutants combined (203 and 228, respectively) (Fig. 2C and Supplemental Fig. S4A). PCA of myoblast and early myotube stages revealed that DEL_SM clones and FSHD patient myocytes cluster together (Fig. 2A). While PC1 (49.2% variance) largely separates myoblast and differentiated myotube stages, there is a significant clustering of patient and DEL_SM myotubes comparing to the control and single mutants, and two clusters are further separated in PC2 (14.0% variance) (Fig. 2A). The top DEGs in PC1 include myogenesis genes that are downregulated (Supplemental Table. S2). Box plots show that albeit at varying degrees, these genes are commonly downregulated in all mutants at early myotube stage (Fig. 2B, top; Supplemental Table S1). Thus, all three types of mutations (DEL, SM and DEL_SM) impede expression of muscle genes. The top mis-expressed genes in PC2 include a subset of DUX4 target genes that are significantly upregulated in patient and DEL_SM, but not in single mutant, early myotubes (Fig. 2B, bottom; Supplemental Fig. S4B).

**Figure 2.**
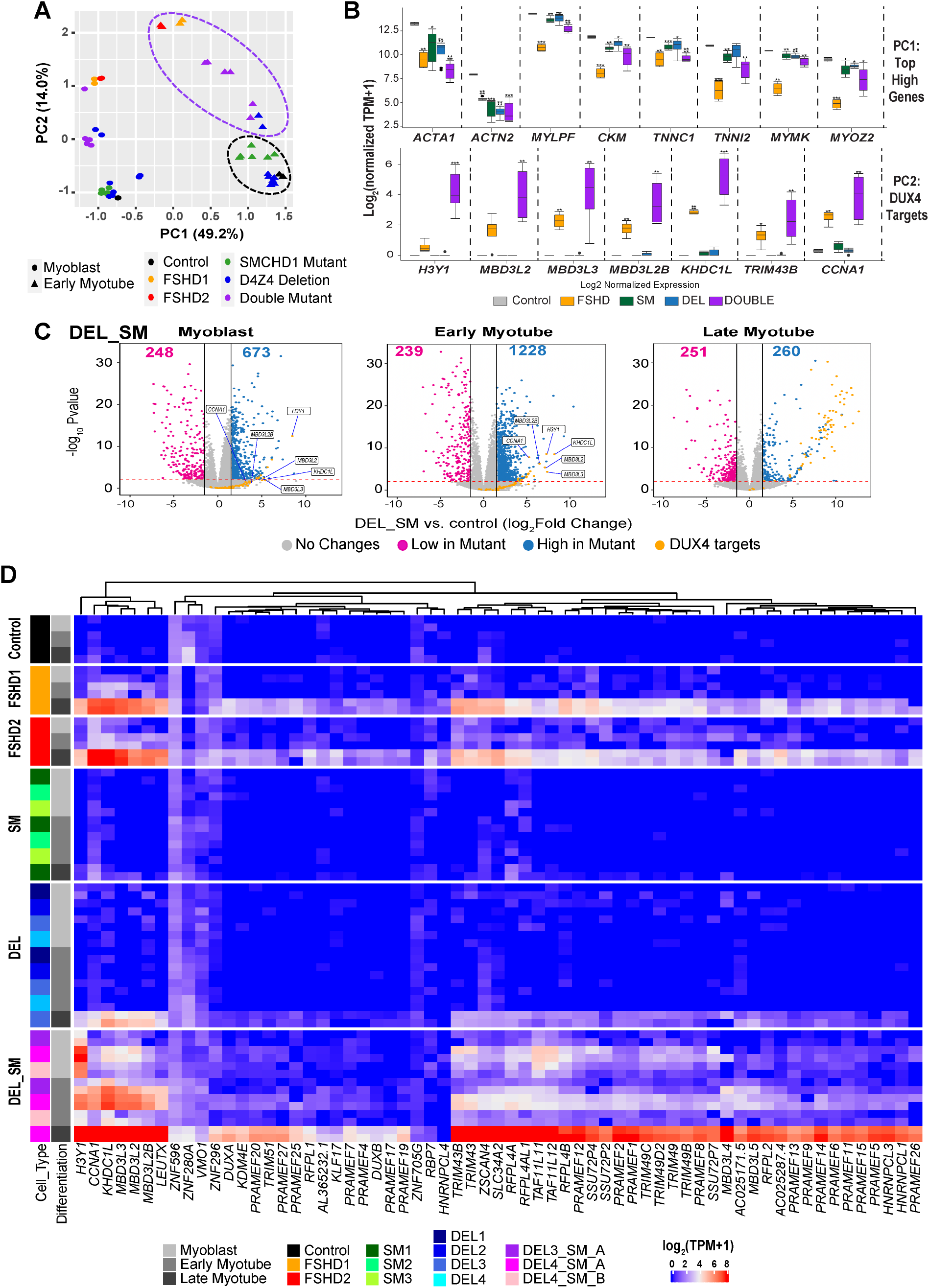
Double mutants closely recapitulate patient cells. **A.** PCA analysis of myoblasts and early myotubes across all the cell types and clones. Top genes for each component are included in the Supplemental Table S2. Differentiation days are indicated by shapes and cell types are indicated by colors according to the label legend. **B.** Expression comparison of selected genes from PC1 and PC2 from **(A)**. Top: box plots of the selected top high genes expression of PC1 in control, FSHD patients and 3 types of mutant early myotubes. Bottom: box plots of 7 DUX4 targets expression from the top 500 high genes of PC2. Expression values are in log2 (normalized TPM +1). Significant values were calculated by Wilcoxon t-test (**** *P* < 0.0001, *** *P* < 0.001, ** *P* < 0.01, * *P* < 0.05). **C.** Volcano plots of significance (log_10_ *P*-value) and log_2_ fold change of double mutants (DEL_SM) compared to control at myoblast, early and late myotube stages. Significantly upregulated (blue) and downregulated (pink) genes and DUX4 targets (orange) are shown. DUX4 target genes in PC2 **(B)** are indicated in myoblasts and early myotubes. **D.** Hierarchical heatmap of DUX4 target gene expression. A total of 64 target genes were selected based on previous studies (38, 42). Expression values are in normalized TPM and log transformed. Grey shades indicate differentiation and colors indicate cell types.

Expression of 64 representative DUX4 target genes (38, 42) increased in DEL_SM during differentiation and became predominant in late myotubes (Fig. 2C, orange). Hierarchically clustered heatmaps in three types of mutant clones compared to isogenic control and FSHD patient myocytes indicated that DUX4 target gene expression is strongest at late myotube stage in DEL, DEL_SM and patient cells (Fig. 2D). In SM mutants, however, DUX4 target expression was over all very low throughout differentiation (Fig. 2D). Consistent with the RT-qPCR analyses (Fig. 1E and I), target gene expression is highly prominent in DEL_SM mutants, which is clear even in myoblast and early myotube stages (Fig. 2D; Supplemental Figs. S4B and S5A). This is in a stark contrast to DEL mutants, in which target gene expression is relatively weak in early stages (Fig. 2D; Supplemental Fig. S4B and S5A). These results further highlight the synergy between D4Z4 contraction and SMCHD1 KO.

### Characteristics of non-DUX4 target DEGs are recapitulated in mutant cells

We performed Gene Ontology (GO) enrichment analysis on the DEGs from DEL_SM for three stages of differentiation (myoblasts, early and late myotubes) (Fig. 3A and B; Supplemental Table S3). While similar GO terms were enriched for downregulated genes at all three stages, we found largely distinct GO term enrichment for upregulated genes at each stage, suggesting that upregulation of DEGs are closely linked to differentiation stages of myocytes (Fig. 3A and B).

**Figure 3.**
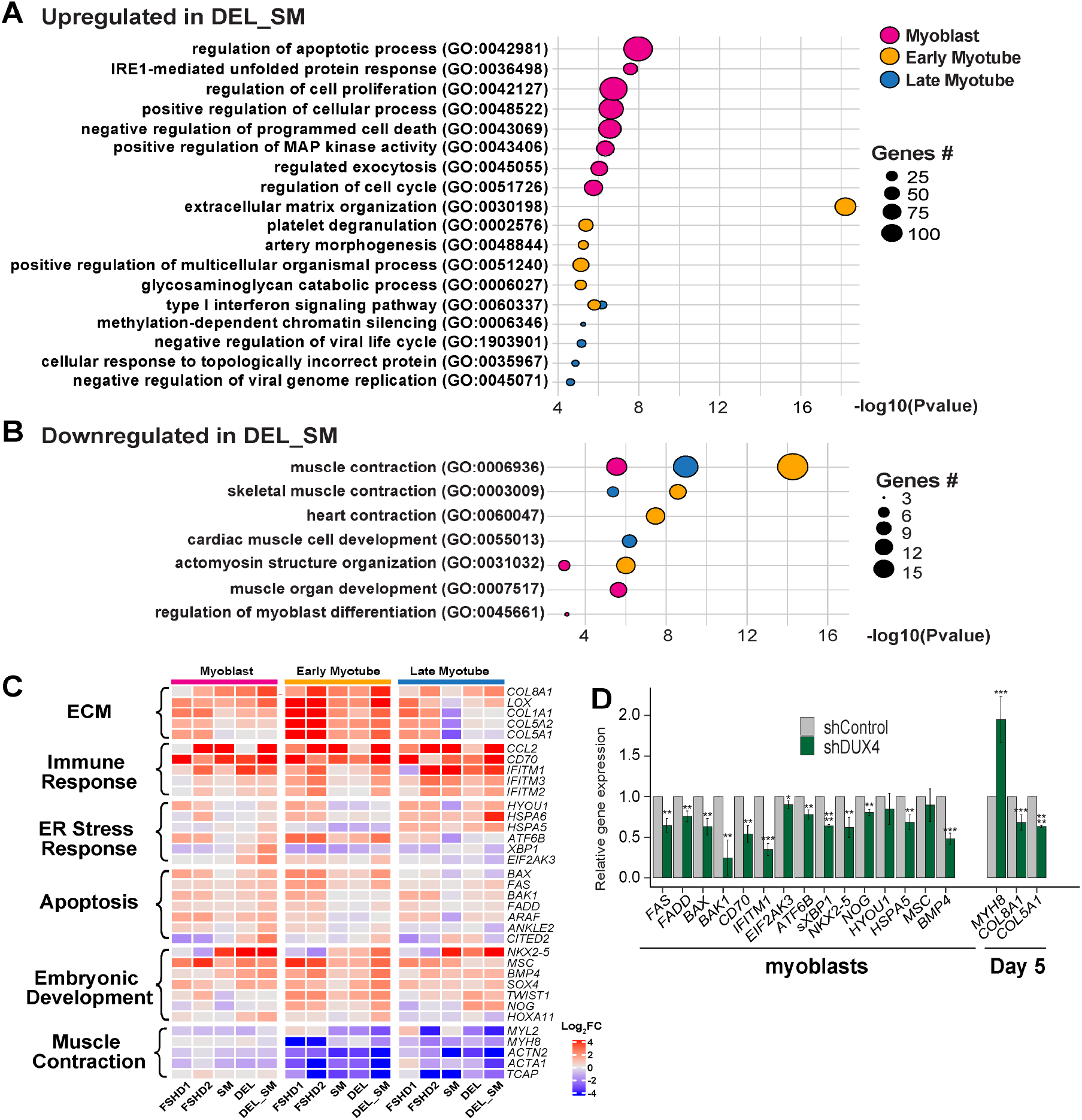
Ontology analyses of common and distinct gene expression in patient and double mutant cells. **A.** The bubble plot shows gene ontology enrichment analysis of upregulated genes in the double mutants in myoblasts (pink), early (yellow) and late (blue) myotubes. The plot shows the selected top terms for each differentiation stage. X-axis displays log_10_ *P*-value and bubble size indicates number of genes in each term as indicated. **B**. Similar to **(A)**, bubble plot for downregulated genes. **C.** Heatmap of log_2_ fold change expression for the selective genes in FSHD patients and mutants and their related pathway. **D.** The expression level of selected genes from Figure 3C in double mutant myoblast (DEL4_SM_A) transduced with lentivirus carrying shControl or shDUX4. Real-time RT-qPCRs were performed for three biological replicates for each sample. Data are presented as mean ± SD; **p<0.01, ***p<0.001, by one-tailed student’s t-test. Results presented as fold difference compared to shControl sample.

Notably, we observed prominent upregulation of genes related to extracellular matrix (ECM), immune response, ER stress, apoptosis, and embryonic genes as well as down regulation of muscle-related genes across patient and mutant cells (Fig. 3C). Similar gene expression changes have been reported in FSHD patient myocytes (16, 35, 45) and some in the recombinant DUX4 overexpression study (10). Our results indicate that D4Z4 and *SMCHD1* mutations are sufficient to recapitulate this patient gene expression phenotype. Importantly, DUX4 depletion reversed these changes of representative genes, strongly suggesting that most of these changes are triggered by the mutation-induced DUX4 expression (Fig. 3D). However, DUX4 may regulate these genes indirectly as the depletion effect was not as robust as that on defined DUX4 target genes (Fig. 1F).

Both positive and negative regulators of apoptosis are upregulated especially in patient, DEL and DEL_SM myoblasts and early myotubes (e.g., *BAX, BAK1, FAS* and *FADD* for pro-apoptosis, and *CITED2, ANKLE2*, and *ARAF* for cell survival) (Supplemental Table S3). Unexpectedly, these gene expression changes appear to taper off in late myotubes when DUX4 target genes are most highly expressed (Figs. 2C and D, 3C). This is consistent with our previous observations that no significant evidence for apoptosis/necrosis was detected in FSHD patient cells (data not shown) (37, 38). These results argue against the cytotoxic fate of DUX4-activated cells.

### DEL mutations diminish D4Z4 heterochromatin, which is exacerbated by H3K9me3 reduction induced by SMCHD1 KO

Previously it was shown that disruption of heterochromatin integrity at the *DUX4* promoter region in the D4Z4 repeat is linked to DUX4 de-repression in both FSHD1 and FSHD2 myoblasts (24, 25). To interrogate the status of H3K9me3 and DNA methylation at the DUX4 promoter region, we performed ChIP-qPCR and MeDIP, respectively, in several clones each of mutant myoblasts (Fig. 4A and B). The reduction in H3K9me3 was observed in both DEL and SM, and was more substantial and consistent in DEL_SM (Fig. 4A). Thus, the loss of *SMCHD1* affects H3K9me3 at the *DUX4* promoter, and further enhances H3K9me3 reduction in D4Z4 mutants, correlating with their synergy on DUX4 and target gene expression (Figs. 2 and 4A). We previously showed that SMCHD1 binding to D4Z4 is H3K9me3-dependent (25). Thus, combined, our results uncover a positive feedback relationship between SMCHD1 and H3K9me3 at D4Z4.

**Figure 4.**
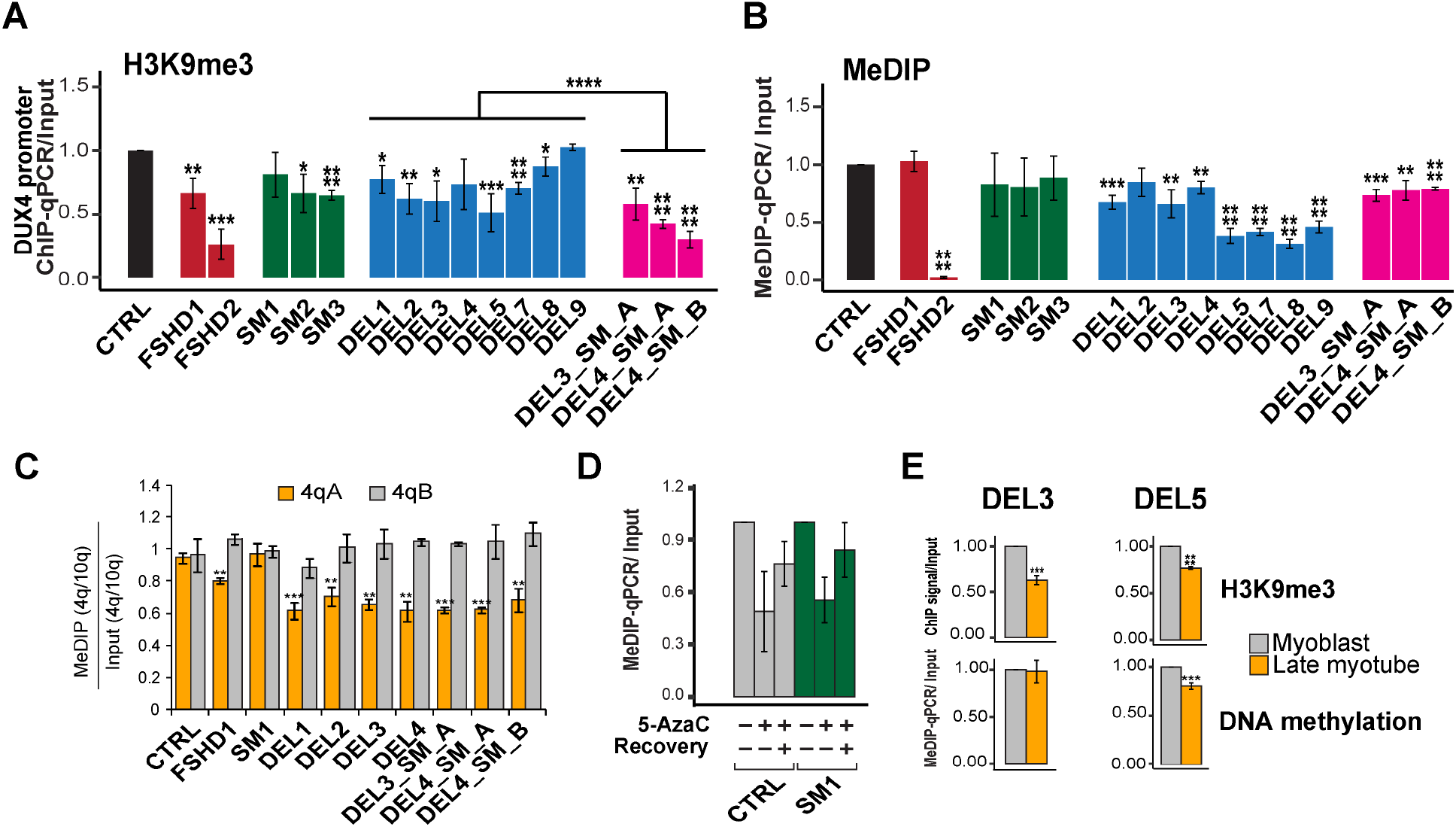
Heterochromatin changes in mutant cells. **A.** H3K9me3 ChIP-qPCR analysis of the *DUX4* promoter region in FSHD1, FSHD2 and mutant myoblasts. Reduction of H3K9me3 is enhanced in double mutant cells. For both **(A)**and **(B)**, signals were normalized to input. Significant comparisons to the control are shown with the asterisks calculated by student’s t-test (**** *P* < 0.0001, *** *P* < 0.001, ** *P* < 0.01, * *P* < 0.05). **B.** DNA methylation from MeDIP analysis. No reduction of DNA methylation was observed in SMCHD1 only and no additional effect in double mutant myoblasts. **C.** Comparison of DNA methylation levels at 4qA, 4qB, and 10q D4Z4 regions among the control, FSHD, and mutant cells. The MeDIP and input samples from **(B)**were amplified by using 4q/10q-D4Z4 specific PCR primers. The PCR products were sequenced and the 4qA, 4qB, and 10q D4Z4 specific sequence reads were analyzed. The 4qA(orange)/10q and 4qB(grey)/10q ratios of MeDIP were normalized with that of input. The data indicated relatively lower methylation at 4qA (but not 4qB) D4Z4 regions of FSHD1, D4Z4 deletion mutants and double mutants. *P*-values for significant differences versus the control sample are shown. **D.** The effect of 5AzaC treatment on SMCHD1 mutants on DNA methylation. The control or SM1 cells were treated with 5AzaC for 24 hours and allowed for 48 hours of recovery before harvesting for MeDIP-qPCR. No significant differences were observed. **E.** H3K9me3 ChIP-qPCR and MeDIP analysis at the DUX4 promoter region were performed to compare between Day 0 and Day 14 of DEL3 or Day 12 of DEL5 mutants. (**** *P* < 0.0001, *** *P* < 0.001).

In contrast to FSHD2 patient cells carrying *SMCHD1* mutations, loss of *SMCHD1* in our cell lines did not affect DNA methylation (Fig. 4B). Thus, a somatic mutation of *SMCHD1* by itself does not recapitulate the DNA methylation changes observed in FSHD2 myoblasts. The relatively subtle changes of MeDIP signals in our D4Z4 mutant cells (and no apparent change in FSHD1 patient cells) may be due to the fact that PCR primers do not distinguish 4q and 10q D4Z4. While DNA hypomethylation was shown to occur at both 4q and 10q alleles in FSHD2 cells, the changes occurred only in the contracted 4qA D4Z4 in FSHD1 cells (21–23). To distinguish different alleles, we separated reads specific to 4qA, 4qB and 10q D4Z4 based on their SNPs (24) and analyzed the read count ratio of 4q/10q in MeDIP compared to input genomic DNA at the *DUX4* promoter region (Fig. 4C). This allowed us to observe a significant decrease of MeDIP signals in FSHD1, DEL and DEL_SM mutants at 4qA D4Z4 over 10q, but not at 4qB over 10q (Fig. 4C). Importantly, we observed no significant difference between control and SM1 mutant cells as well as between DEL and DEL_SM cells (Fig.4C). These results suggest a negligible contribution of *SMCHD1* KO to the maintenance of D4Z4 DNA methylation in adult myocytes.

An earlier study suggested that SMCHD1 is important for de novo methylation of D4Z4 repeats (43). Therefore, we compared recovery of DNA methylation in control- or SM1 mutant myoblasts after treatment with the DNA methylation inhibitor, 5AzaC. Following treatment with 5AzaC for 48 hours, cells were allowed to recover for 48 hours. MeDIP revealed no differences in recovery between the control and *SMCHD1* mutant cells (Fig. 4D). Taken together, our results indicate that SMCHD1 is not required for either the maintenance or the re-establishment of DNA methylation at the DUX4 promoter in adult myoblasts. It remains possible that *SMCHD1* must be mutated earlier in development to have an effect on DNA methylation at D4Z4, or there may be an additional mechanism/modifier gene involved in FSHD2.

At late myotube stages in DEL mutants, we observed a further reduction in DNA methylation and/or H3K9me3 at the *DUX4* promoter, accompanied by more efficient expression of *DUX4* and target genes (Figs. 2D and 4E). In conclusion, our results indicate that the degree of loss of heterochromatin caused by FSHD mutations correlates with the degree of activation of the DUX4 network and severity of the disease.

### Inhibition of DNA methylation boosts expression of the DUX4fl network in mutant cells

One major difference between our somatic mutant cells and FSHD2 cells that were tested was the level of DNA methylation (Fig. 4B). To examine the role of DNA methylation, we treated control and mutant cells with 5AzaC. We observed robust induction of DUX4 target genes in DEL and DEL_SM mutant, but not in control cells (Fig. 5A; Supplemental Fig. S5B). Upregulation is more prominent in early myotubes than at myoblast stages (Fig. 5A). Although DUX4 target genes were significantly upregulated in SM myotubes by 5AzaC, they were induced over 100-fold higher in DEL myotubes (Fig. 5B and C). Importantly, in both cases, upregulation was DUX4-dependent (Fig. 5D; Supplemental Fig. S6A). Because 5AzaC also inhibits RNA methylation, we tested 5AzadC, which only gets incorporated into DNA, and obtained the same results (Supplemental Fig. S6B). Moreover, MeDIP indicates that both 5AzaC and 5AzadC caused a comparable reduction in DNA methylation at D4Z4 (Supplemental Fig. S6C). We conclude that inhibition of DNA methylation leads to an induction of the DUX4 gene network.

**Figure 5.**
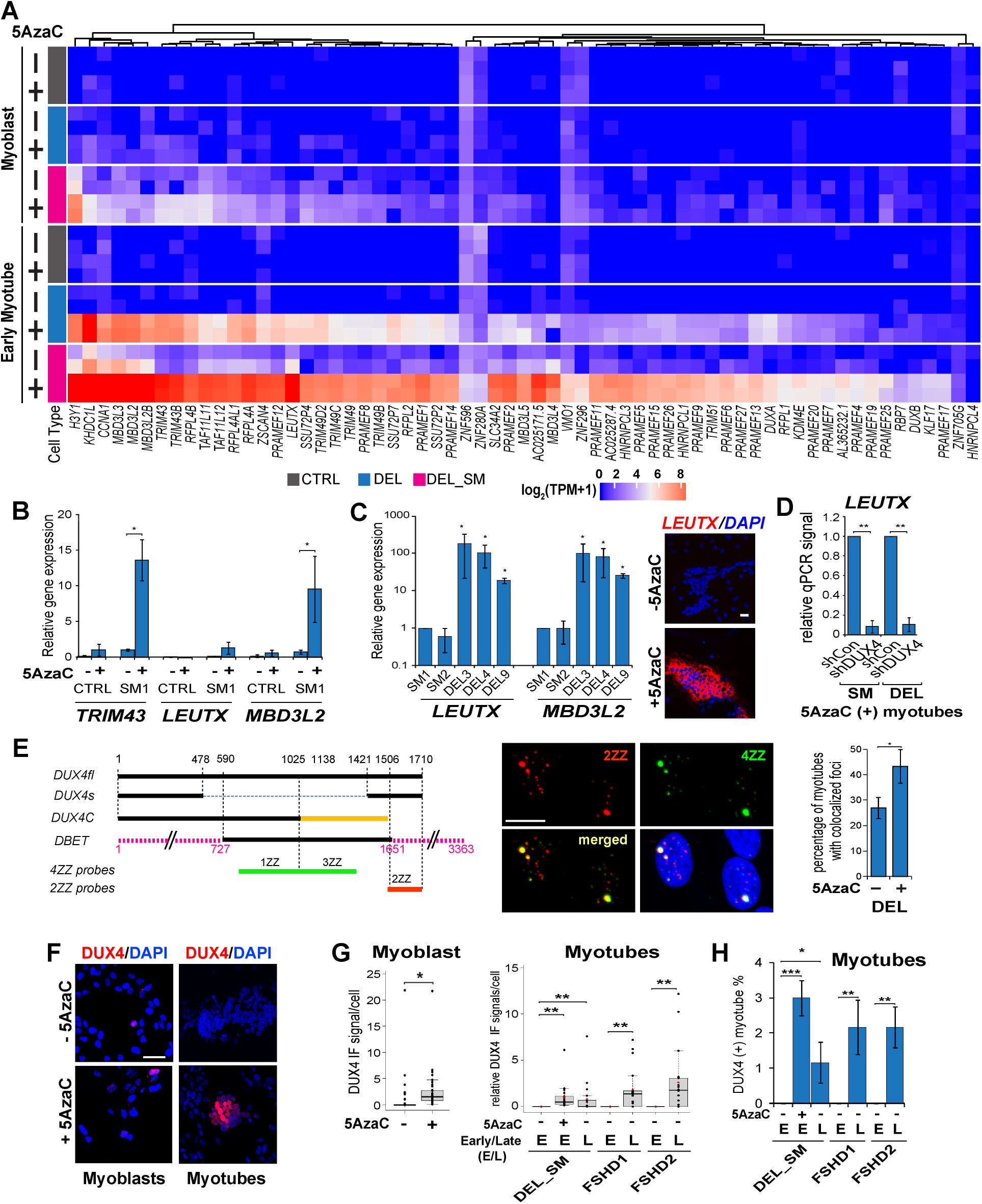
Inhibition of DNA methylation increases DUX4fl RNA and protein and robustly upregulates target genes in mutant cells. **A**. Hierarchical heatmap of DUX4 target gene expression for control, DEL3, and DEL3_SM_A mutants with or without 5AzaC at myoblast and early myotube. A total of 64 target genes were selected based on previous studies (38, 42). Expression values are in normalized TPM and log transformed. **B.** DUX4 target genes were greatly affected by 5AzaC treatment in mutant cells. Control and SM1 myoblasts were treated with or without 5AzaC for 48 hrs. Then 5AzaC was removed from the media, and differentiation was induced. 4 days later, RT-qPCR of DUX4 target genes were performed. The gene expression data were normalized to *GAPDH* level in each sample, and then normalized to the *LEUTX* value of 5AzaC treated SM1. Data are presented as mean ± SD; *p<0.05, by one-tailed student’s t-test. **C.** Comparison of DUX4 target genes level between early myotubes of 5AzaC treated SM and DEL mutants. SM and DEL myoblasts were treated with 5AzaC for 48 hrs right before differentiation. At day 5 of differentiation, the mRNA expression level of DUX4 target genes was assessed by real-time RT-PCR, relative to SM1. Data are presented as mean ± SD; *p<0.05, by one-tailed student’s t-test. Representative images of in situ detection of *LEUTX* RNA (red) with or without 5AzaC treatment are shown on the right (blue: DAPI). Scale bar 10 μm. **D.** DUX4 depletion inhibited DUX4 target gene upregulation induced by 5AzaC treatment in mutant cells. SM1 and DEL3 cells were treated with 5AzaC and induced differentiation same as **(B)**. During 5AzaC treatment, cells were infected with lentivirus containing shCTRL or shDUX4. For each cell line, LEUTX expression level after DUX4 depletion was shown as fold difference compared to the control. Data are presented as mean ± SD; **p<0.01, by one-tailed student’s t-test. **E.** 5AzaC facilitated DUX4fl expression in DEL3 early myotubes. Left: the schematic diagrams of mRNA transcripts for *DUX4fl*, the *DUX4s* isoform and *DUX4* homologs (*DUX4c* and *DBET*), the black regions, which represent >99% homology to *DUX4fl*, could be detected by corresponding *DUX4* 4ZZ probes or 2ZZ probes, but not by both. Therefore, the overlapping signals from 4ZZ and 2ZZ probes represent the *DUX4fl* transcripts. Middle panel, example images of the RNAScope results of the 4ZZ probes (green), 2ZZ probes (red) and the overlapping foci (yellow). DAPI is in blue. Scale bar = 10μm. Right, 5AzaC treatment increased the percentage of myotubes with overlapping foci of 4ZZ and 2ZZ probes. Data from 3 independent experiments are presented as means ± SD. *p<0.05, by one-tailed student’s t-test. **F.** Examples of DUX4 protein expression in double mutant myoblasts and myotubes. Immunofluorescence for DUX4 on DEL4_SM_A cells after 5 days of differentiation. Nuclei were counterstained with DAPI (blue). Scale bar 50 μm. **G.** Quantification of DUX4 protein expression with and without 5AzaC in double mutant myoblasts (left) and early myotubes (right). Mutant late myotubes and FSHD1 and FSHD2 patient early and late myotubes are shown for comparison. The DUX4 integrated density values in myoblasts/myotubes were measured using ImageJ software (37). Top 3% values in each group were used for graph and data analysis. All the data were normalized to the corresponding mean value of the 5AzaC-treated DEL_SM samples. *p<0.05, ** p<0.01, ***p<0.001. (Totally 600 myotubes or 1200 myoblasts were observed in each group). **H.** Replotting the data in (G, right panel) for the frequency of DUX4 IF staining positive myotubes. *p<0.05, ** p<0.01, ***p<0.001.

To specifically detect the *DUX4fl* transcript that can be translated into the protein, we split our RNAScope probes to 4ZZ specific to the middle region and 2ZZ specific to the 3’end of the DUX4fl transcript (Fig. 5E). Therefore, colocalization of the two probes should reflect the presence of DUX4fl. Using this strategy, we were able to confirm that the prominent nuclear foci of *DUX4* RNA previously detected (37) represents *DUX4fl* transcripts (Fig. 5E). 5AzaC treatment indeed increased the colocalized signals of *DUX4fl* in DEL mutant myotubes (Fig. 5E, right). Consistently, DUX4 protein expression at the early myotube stage is significantly induced by 5AzaC treatment to a level that is comparable to those at late myotube stages of DEL_SM and patient cells (Fig. 5F-H). Collectively, these findings indicate that heterochromatin disruption stimulates target gene induction through upregulation of DUX4fl expression in mutant cells. Moreover, we found that contraction of D4Z4 makes this locus highly sensitive to additional heterochromatin destabilization, emphasizing a potential role for epigenetic modifiers in the disease penetrance and severity in FSHD1.

### Coherent feedforward loop of early and late DUX4 target genes

*LEUTX* is a DUX4 target gene that encodes for a transcription factor critical for zygotic genome activation in early development (46). We observed significant increase of *LEUTX* transcript and protein signals in early mutant myotubes after 5AzaC treatment (Figs. 5C and 6A, respectively). Interestingly, however, LEUTX induction in myoblasts is very weak (Fig. 6A). This is in contrast to the significant induction of another DUX4 target H3.X/Y in myoblasts by 5AzaC (Fig. 6A). We found that genes, such as *H3Y1, MBD3L2, KHDC1L*, are among the top DUX4 target genes activated, and are efficiently expressed and can be stimulated by 5AzaC in the myoblast stage in DEL_SM cells (Figs. 2D and 6C; Supplemental Fig. S7). In contrast, *LEUTX, DUXA, RFPL1* and *KLF17*, are expressed only at a low level and fail to get stimulated by 5AzaC in the myoblast stage and are induced much more efficiently after cells were differentiated into myotubes (Fig. 6C; Supplemental Fig. S7). These results reveal that there are two classes of DUX4 targets: early response genes and differentiation-dependent late target genes.

**Figure 6.**
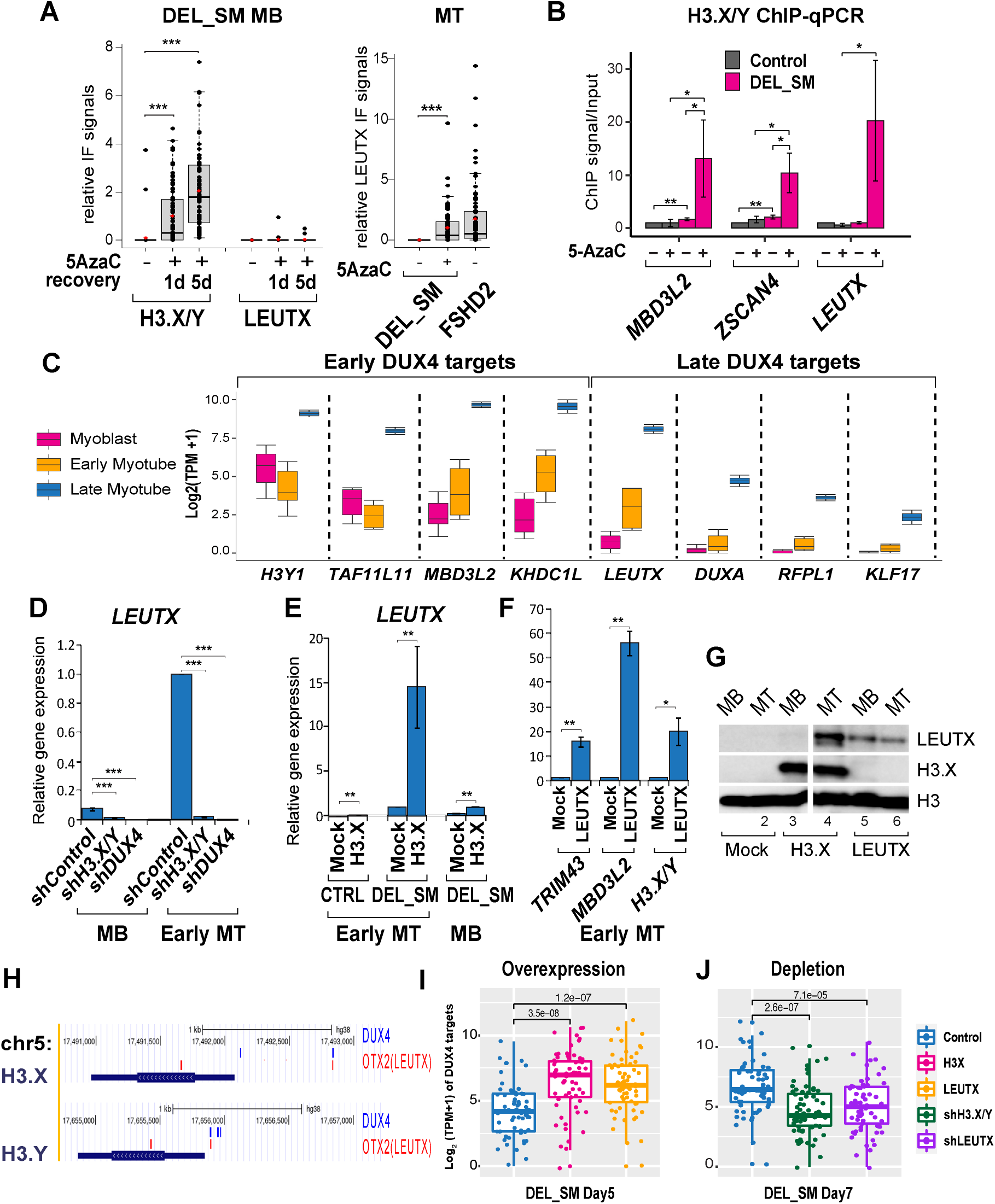
Identification of early and late DUX4 target genes that are differentially sensitive to myotube differentiation. **A.** Double mutant cells were treated with or without 5AzaC as indicated. IF signals of H3.X/Y and LEUTX in myoblasts (left panel) and LEUTX in early myotubes (right panel) were quantified as integrated intensity in each myoblast/myotube using ImageJ software. FSHD2 patient early myotubes are shown for comparison. The DUX4 integrated density values in each myoblast/myotube were measured using ImageJ software. Based on the highest positive myoblasts/myotubes number of all, same number of values in each group were used for graph and data analysis. All the data were normalized to the corresponding mean value of the 5AzaC treated samples (release day1 for the myoblasts). Red dots represent mean values. (N=300 myotubes or 1000 for myoblasts). ***p<0.001, by one-tailed student’s t-test. **B.** Incorporation of H3.X/Y into DUX4 targets in control and double mutant at Day 4 with or without 5AzaC. Cells were treated with 5AzaC for 48 hours before differentiation. Significant incorporation of H3.X/Y is shown by the asterisks with the indicated comparisons. **C.** Box plots of representative early and late DUX4 target gene expression in double mutant cells were shown to compare myoblast, early and late myotube stages as indicated. Expression values are in log2 normalized TPM. **D.** The expression level of DUX4 target gene LEUTX in double mutant DEL4_SM_A transduced with lentivirus carrying shControl, shH3.X/Y or shDUX4. Real-time RT-PCRs were carried out before or 5 days after the induction of differentiation. Three biological replicates for each sample were performed. Data are presented as mean ± SD; **p<0.01, ***p<0.001, by one-tailed student’s t-test. Results presented as fold difference compared to shControl differentiated sample. **E.** Control and DEL4_SM_A myoblasts were transduced with a lentiviral empty vector or a lentiviral vector expressing H3. X. Differentiation was induced at 48 hours after transduction. For myoblasts or early myotubes as indicated, the mRNA expression level of the downstream target genes was assessed by real-time RT-qPCR. Data are presented as mean ± SD; *p<0.05, **p<0.01, ***p<0.001, by one-tailed student’s t-test. Results presented as fold difference compared to empty vector infected double mutant cells. **F.** Similar experiments as in **(E)**, but DEL4_SM_A myoblasts were transduced with a lentiviral vector expressing LEUTX. **G.** Overexpression of H3.X and LEUTX in MB or MT was assessed by western blot. Pan histone H3 antibody was used as control as indicated. Lanes 1 and 2: mock transfection. Lanes 3 and 4: H3.X OE. Lanes 5 and 6: LEUTX OE. The endogenous LEUTX is upregulated in H3.X OE myotubes (lane 4). **H.** TF binding motifs at the promoter of H3.X/Y. Binding motifs for DUX4 and the putative LEUTX motif (OTX2) within 1 kb upstream and 0.5 kb downstream of the transcription start site for H3.X/Y were identified by using the MoLoTool provided in HOCOMOCO v11, with *P*-values less than or equal to 0.001. Visualization was done on the UCSC genome browser using GENCODE v36 for the H3.X/Y genes model. **I.** The effects of control, H3.X/Y or LEUTX overexpression (as in **E-G**) on 64 DUX4 target genes in DEL_SM myotubes Day 5 are assessed by RNA-seq and displayed in box plots. *P*-values are calculated using Wilcoxon t-test indicated at the top. **J.** Similar analysis was performed with H3.X/Y or LEUTX shRNA depletion compared to the same control as in **(I)** on DUX4 target gene expression in DEL_SM myotubes Day 7.

*Histone H3Y1 (H3.Y*) is one of the highest induced DUX4 targets in mutant cells (Fig. 2D). H3.X/Y was found to be incorporated in the target gene regions and increased their expression (47). Indeed, H3.X/Y binding to *MBD3L2, ZSCAN4* and *LEUTX* gene regions was all induced in 5AzaC-treated DEL_SM, but not control myotubes, providing an additional mechanism for increased target gene expression by 5AzaC (Fig. 6B). Consistently, shRNA depletion of H3.X/Y effectively blocks *LEUTX* activation in DEL_SM myotubes (Fig. 6D). The effect is nearly comparable to DUX4 depletion, emphasizing the strong reliance of DUX4 target gene expression on H3.X/Y (Fig. 6D). Conversely, overexpression of H3.X significantly stimulated *LEUTX* expression in DEL_SM cells (Fig. 6E). However, consistent with the notion that *LEUTX* is a differentiation-dependent late target gene, induction of LEUTX mRNA and protein is much more robust in myotubes than in myoblasts, despite comparable H3.X/Y overexpression in both cells (Fig. 6E and G).

Overexpression of LEUTX also stimulated *H3.X/Y* (as well as *MDB3L2* and *TRIM43*), indicating a positive feedback loop (Fig. 6F and G). Indeed, H3.X and H3.Y gene promoters both contain LEUTX (OTX2) binding motifs, raising the possibility that LEUTX directly upregulates *H3.X/Y* genes (Fig. 6H). Intriguingly, genome-wide analyses of overexpression and depletion of H3.X/Y and LEUTX by RNA-seq reveal that both of them globally affect DUX4 target gene expression (Figs. 6I and J). Taken together, the results indicate that the differentiation-independent early DUX4 target H3.X/Y gets incorporated and sets the epigenetic stage for upregulation of other DUX4 target genes, including the differentiation-dependent LEUTX. Once expressed, LEUTX further stimulates expression of H3.X/Y as well as other DUX4 target genes to fortify the DUX4-triggered gene network. Taken together, our results suggest that the DUX4 gene network is composed of a coherent feedforward portion with DUX4 working together with H3.X/Y to drive other target genes as well as a positive feedback loop between H3.X/Y and LEUTX (and possibly other TFs) that is subsequently activated and locks in target gene expression (Fig. 7B).

**Figure 7.**
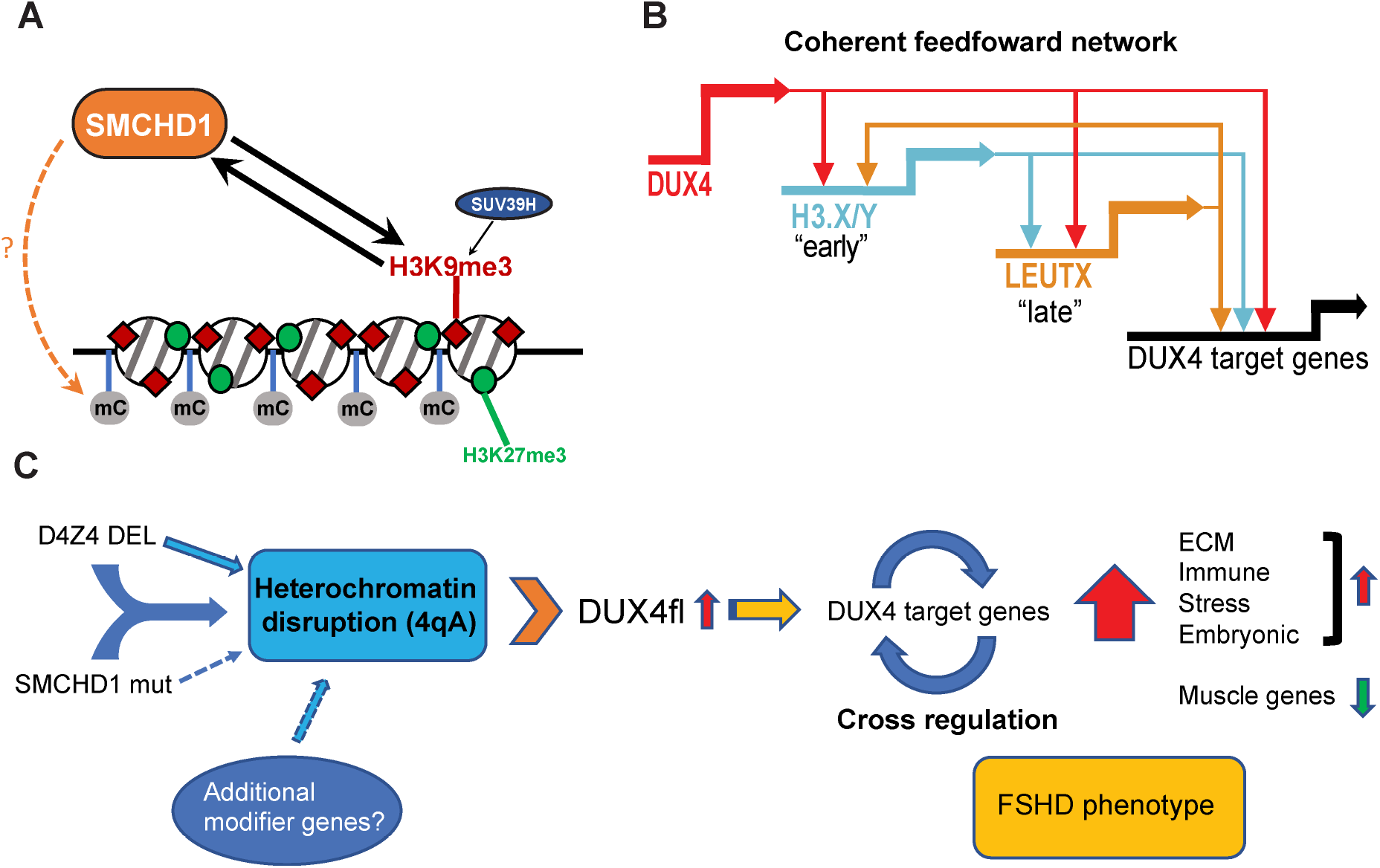
**A.** Positive feedback loop between H3K9me3 and SMCHD1. SMCHD1 interacts with D4Z4 chromatin in an H3K9me3-dependent manner (25) and also maintain H3K9me3. **B.** Coherent feedforward mechanism of DUX4 and target gene expression. While DUX4 is critical for the initial activation of its target genes, the early target H3.X/Y expression is essential for efficient expression of other downstream target genes, including the late target TF, LEUTX. LEUTX in turn promotes further expression of H3.X/Y. H3.X/Y as well as LEUTX (and possibly other DUX4 target TFs) contribute significantly to the expression of other DUX4 target genes. **C.** Two key processes in FSHD pathogenesis. D4Z4 heterochromatin disruption induced synergistically by D4Z4 and SMCHD1 mutations (and possibly other epigenetic modifiers) enables stabilization and enhancement of DUX4fl expression. Once activated by DUX4, DUX4 target genes undergo cross-regulation contributing to the establishment of the FSHD gene expression phenotype.

## Discussion

In the present study, we investigated the effects of genetic FSHD mutations in healthy isogenic human skeletal myoblast cells. Particularly, we compared effects of D4Z4 contraction or loss of *SMCHD1*, either alone or combined. We found that, although loss of *SMCHD1* by itself has only minor consequences, this mutation strongly synergizes with D4Z4 contraction, resulting in upregulation of the DUX4fl target gene network. These results support the notion of SMCHD1 mutations as modifiers that enhance FSHD1 severity (7, 8). Interestingly, SMCHD1 mutation affected H3K9me3 but not DNA methylation (Fig. 7A). Our analyses uncovered two key processes in the establishment of the robust FSHD disease phenotype: (1) disruption of heterochromatin at D4Z4 repeats and (2) coherent feedforward loop of the DUX4 target gene expression (Fig. 7B and C). Our mutant cell models closely recapitulate the patient cell phenotype and display sensitivity to disease modifiers. Importantly, the analysis of late myotubes allowed us to separate robust DUX4 target gene activation from cytotoxicity/apoptosis. Our results suggest that transcriptional activation of the DUX4 target gene network, rather than shortterm cytotoxic effects, is relevant for FSHD disease progression.

Our CRISPR mutation using gRNA specific to 4q D4Z4 repeat upstream of the *DUX4* transcription start site has led to repeat contraction as initially assessed by PFGE. A similar CRISPR approach was used to remove telomere repeats in human neuroblastoma cells (48). We found that despite some repeat inversion or, CRISPR plasmid insertion to D4Z4 in one set of DEL mutants, the last copy of 4qA D4Z4 remained intact enabling the expression of *DUX4fl*. Perhaps it is not surprising that repetitive sequences can be recombinogenic, which makes it important to assess repeat alteration by direct sequencing of genomic DNA and transcript RNA using nanopore long-read sequencing.

We previously demonstrated that D4Z4 heterochromatin is marked by H3K9me3, which is required for the recruitment of HP1γ, cohesin and SMCHD1 (24, 25). Reduction of H3K9me3 leads to decrease of SMCHD1 binding, accompanied by increased expression of DUX4 (25). In the currently study, albeit more variable than in DEL mutants, we found that *SMCHD1* mutation reduced H3K9me3 and increased DUX4 target gene expression in a modest, but statistically significant manner, suggesting the positive feedback loop between SMCHD1 and H3K9me3 contributing to *DUX4* suppression. Interestingly, reduction of H3K9me3 was also observed in D4Z4 transgenic mice crossed with smchd1 mutant mice (49). Furthermore, a recent study showed that restoration of SMCHD1 expression in FSHD2 iPSCs increased the level of H3K9me3 and HP1γ at D4Z4 chromatin (50). SMCHD1 is recruited to and compacts human inactive X chromosome in part through interaction with HP1-bound H3K9me3 chromatin via HP1-binding protein HBiX1 (LRIF1) (51, 52). Suggestively, a homozygous mutation of *LRIF1* has also been linked to FSHD2 (53). Thus, disruption of the observed positive feedback loop between SMCHD1 and H3K9me3 at D4Z4 may be critical for FSHD pathogenesis.

Smchd1 was originally identified to play a role in the maintenance of DNA methylation at CpG islands and inactive X chromosome (27–29). Since FSHD2 cells (with *SMCHD1* mutations) tend to exhibit a strong DNA hypomethylation phenotype (21) (and this study), it was speculated that SMCHD1 mutation results in loss of DNA methylation leading to DUX4 upregulation. A recent study showed that Smchd1 binds and antagonizes Tet enzymes (54). Consequently, loss of *Smchd1* in mouse ES cells leads to reduced DNA methylation and upregulation of *Dux*, a functional homolog of human *DUX4* (54). Our results, however, suggest that such a DNA methylation-dependent mechanism does not appear to operate in human myocytes. We cannot formally exclude the possibility that there is a narrow developmental window in which haploinsufficiency of SMCHD1 effectively blocks the initial establishment of DNA methylation at D4Z4. Indeed, it was shown that D4Z4 remethylation during reprogramming cannot take place in SMCHD1-mutated FSHD2 iPSCs, suggesting that SMCHD1 is involved in the initial establishment of D4Z4 DNA methylation earlier in development (43). However, since that study were done using FSHD2 iPSCs rather than SMCHD1 knockout iPSCs, possible contributions of additional modifier gene(s) in FSHD2 iPSCs cannot be excluded. Interestingly, *SMCHD1* gene correction in FSHD2 iPSCs failed to increase DNA methylation (50). The fact that *SMCHD1* mutation did not affect D4Z4 DNA methylation, yet cooperated strongly with D4Z4 contraction in activating DUX4 target gene expression in our mutant cells, suggests that SMCHD1 suppresses DUX4 in a DNA methylation-independent manner. This might involve the SMCHD1-H3K9me3 positive feedback loop discussed above. This is consistent with a previous report that SMCHD1 can suppress gene expression in a DNA methylation-independent manner (30). Interestingly, we observed that 5AzaC-induced DNA hypomethylation in *SMCHD1* mutant cells is not sufficient to recapitulate the robust phenotype of FSHD2 patient cells (with *SMCHD1* mutation) used in our study. These observations hint at the existence of additional factors that synergize with *SMCHD1* mutations to generate full FSHD2 pathogenesis (Fig. 7C).

In addition to DUX4 target genes, D4Z4 disruption combined with loss of SMCHD1 in our mutant cells yielded a gene-expression pattern that recapitulated key features of patient myocytes. These include upregulation of genes related to ECM, immune and stress responses as well as embryonic genes, and downregulation of muscle genes (10, 16, 35, 45, 55). Crucially, DUX4 depletion reversed these changes in gene expression, strongly suggesting that they are DUX4-dependent. In contrast to the almost complete suppression of the DUX4 target gene network, this reversal is only partial, suggesting that expression changes of these other pathways are largely indirect downstream effects. Notably, changes in the expression of genes involved in the regulation of apoptosis (both anti- and pro) are more prominent in myoblasts, and appear to taper off later in myotube differentiation when DUX4 target genes are most highly induced. Our results strongly suggest that activation of the DUX4 target gene network is separate from DUX4-induced cell toxicity. Therefore, further analyses of dynamics and consequences of DUX4 and target gene expression during muscle differentiation will be important.

Our results indicate that DUX4 target genes can be sub-divided into early and late genes based on their differential dependency on myotube differentiation. The highly induced early DUX4 targets, histone variants H3.X and/or H3.Y, were previously shown to be incorporated into the DUX4 target gene regions and promote their expression (47). We found that H3.X/Y depletion almost completely inhibits the expression of LEUTX, a late target gene. Interestingly, another differentiation-insensitive early DUX4 target, MBD3L, can disrupt the repressive functions of MBD2 or MBD3, and upregulate *DUX4* expression as a positive feedback regulator (56, 57). Thus, these early DUX4 targets are epigenetic initiators that set the stage to promote activation of the DUX4 gene network. In contrast, late target genes such as LEUTX, are not induced until cells differentiate into myotubes, suggesting an additional differentiation-coupled mechanism of gene regulation. Nevertheless, overexpression or depletion of H3.X/Y as well as LEUTX globally affects DUX4 target expression. LEUTX may directly bind and control these genes and/or indirectly promote their expression through feedback activation of H3.X/Y (and possibly other TFs), as indicated by our study. Taken together, our results strongly suggest that DUX4 triggers the sequential induction of epigenetic regulators and downstream transcription factors to ensure a coherent feed forward effect on the DUX4 gene network (58).

Using CRISPR engineering, we have created *SMCHD1* and/or D4Z4 mutant human skeletal myoblast lines and examined the mutation effects during myocyte differentiation. Our results highlight the fact that FSHD is a heterochromatin abnormality disorder, in which stabilization of otherwise variegated *DUX4fl* expression from contracted D4Z4 allele by chromatin modifier(s) is a critical driver of FSHD pathogenesis. Furthermore, our observations revealed a hierarchy within the DUX4 target genes, highlighting a coherent feed forward mechanism of the DUX4 target gene network activation involving early and late genes, triggered by the expression of the *DUX4fl* from the last D4Z4 repeat. Our results also suggest that in adult myocytes SMCHD1 functions in the context of H3K9me3 rather than regulation of DNA methylation, raising the possibility of additional modifiers in FSHD2. The mutant cells described in the present study will be a valuable resource for further investigation of the molecular mechanism, and serve as a possible platform for therapy development.

## Materials and Methods

### Generation of SMCHD1 knockout mutants in immortalized permissive control myoblast with CRISPR-Cas9

The CRISPR-SMCHD1-1 and 2 constructs used in this study for induction of doublestrand breaks (DSB) at SMCHD1 gene were designed by CRISPRdirect (https://crispr.dbcls.jp/) (Supplemental Figure S2A). ~3×10^5^ Immortalized human control myoblasts were seeded in a 35mm cell culture dish. One day later, the cells were transfected with both 1 μg of CRISPR-SMCHD1-1 and 1 μg of CRISPR-SMCHD1-2 or CRISPR-SMCHD1-1 alone together with 0.5 μg of a puromycin-resistance plasmid using Lipofectamine 3000. The media was changed after 4 hours. The next day the media was replaced with fresh media containing 2 μg/ml puromycin (Sigma-Aldrich) for 3 days. Single cell clones were isolated by FACS sorting into 96 well plates. Ten to 14 days later, genomic DNA of the single cell clones with good proliferation was extracted using a QuickExtract DNA extraction kit (Epicentre). The exon 23-24 region was amplified by a pair of PCR primer (SMCHD1_PCR) to check for genomic deletions on a 1.5% agarose gel. The deletion mutants were confirmed by Sanger sequencing and western blot. SMCHD1 knock out mutants were identified by western blot and pooled amplicon sequencing. Using custom barcoded primers (SMCHD1_seqPCR), pooled amplicons from multiple individuals were sequenced at Genewiz to determine the genomic sequence at the gRNA target site of each cell line. The off-target loci with the highest prediction scores were amplified by a PCR primer pair (SMCHD1_off_target_PCR) and sequenced by Sanger sequencing. For the SMCHD1 mutant cell lines used in this paper, the corresponding off-target sequences (5’-TTTTCAATTTCAGTCAACGA-3’, chr9:+72135246) were not changed.

### Generation of D4Z4 contraction mutants with CRISPR-Cas9

The CRISPR-Cas9 system was used to delete D4Z4 repeat units on 4qA in permissive control myoblasts. Guide RNAs (gRNAs) target a D4Z4 “1-kb” subregion sequence, which excludes the regions that are repeated elsewhere in the genome (59) as well as the DUX4 gene/promoter region to avoid to induce DUX4 mutation (Supplemental Figure S2B). Based on this sequence, gRNAs were designed for 4q D4Z4 using the CRISPR Design Tool (https://zlab.bio/guide-design-resources) with low predicted off-target effects. Two rounds of CRISPR-Cas9 induced D4Z4 repeat array contraction was performed to obtain D4Z4 contraction mutants with satisfactory repeat number. Cas9 (41815, Addgene plasmid) gRNA-D4Z4-1/ 2 (in gRNA expression vector pH082 pU6-gRNA2.0-GFP) and a puromycin-resistance plasmid were cotransfected into parental myoblast as indicated in Supplemental Figure S2B. One mutant with 10 units of 4qA D4Z4 repeat was used as parental cells for the 2nd round of D4Z4 deletion mutant generation. Alternatively, the Alt-R CRISPR-Cas9 genome editing system (IDT) was used for the recombinant Cas9 protein and gRNA delivery for single and double mutations of D4Z4 and SMCHD1 (DEL5, DEL7, DEL8, DEL9 and DEL9_SM, respectively). CRISPR/Cas9 (1081060, Integrated DNA technologies (IDT))/tracrRNA (1073190, IDT)/crRNA(IDT) RNP were delivered to the myoblasts using CRISPRMAX Cas9 Transfection Reagent (CMAX00003, Invitrogen). Transfections were performed as described in the IDT protocol for “Alt-R CRISPR/Cas9 System”. The differentiation efficiency and DUX4 target gene MBD3L2 expression of the single colony cell lines were tested. Based on the results, several cell lines were subjected to PFGE and blot hybridization to confirm the size of D4Z4 regions at 4q and 10q as well as nanopore genomic sequencing (see below).

### Genomic and RNA nanopore long-read sequencing

Genomic Nanopore libraries constructed from genomic DNA using Cas9 Sequencing Kit (SQK-CS9109), Nuclease-free duplex buffer (IDT Cat # 11-01-03-01), and the following Alt-R CRISPR reagent from IDT: tracrRNA (1073190, IDT) resuspended at 100 μM in TE pH 7.5, Cas9 nuclease V3 (1081060, IDT), 3 different S. pyogenes Cas9 Alt-RTM crRNAs (two upstream target sites 5’-CCTATTAAACGTCACGGACA-3’ and 5’-GATACCGACAGCAATAGTCC-3’ and one downstream target site ‘5-AAATCTTCTATAGGATCCAC-3’) resuspended at 100 μM in TE pH 7.5. The Long Fragment Buffer (LFB) from the Cas9 Sequencing Kit (SQK-CS9109) was used during the wash steps. Libraries were loaded on R9.4.1 Flow Cells (FLO-MIN106D) and sequenced on MinION Mk1B instrument using the MinKNOW software. Oxford Nanopore’s base calling software, Guppy version 6.0.1+652ffd1, was run in super accurate (sup) mode with the dna_r9.4.1_450bps_sup.cfg configuration file.

RNA-seq Nanopore libraries were constructed from 200 fmol Illumina libraries using Ligation Sequencing Kit (SQK-LSK110) and NEBNext^®^ Companion Module for Oxford Nanopore Technologies^®^ Ligation Sequencing (E7180S). The Short Fragment Buffer (SFB) from the Ligation Sequencing Kit (SQK-LSK110) was used during the wash steps. 50 fmol sample libraries were loaded on R9.4.1 Flow Cells (FLO-MIN106D) and sequenced on MinION Mk1B instrument using the MinKNOW software. Oxford Nanopore’s base calling software, Guppy version 6.0.1+652ffd1, was run in super accurate (sup) mode with the dna_r9.4.1_450bps_sup.cfg configuration file. Adapters were trimmed from reads with the Porechop software package by adding a custom adapter sequence that includes both the illumina primer and nanopore adapter (top: 5’-AATGTACTTCGTTCAGTTACGTATTGCTAAGCAGTGGTATCAACGCAGAGTAC-3’ and bottom: 5’-GTACTCTGCGTTGATACCACTGCTTAGCAATACGT-3’). Reversed reads were flipped using a custom script.

### Cell culture and differentiation

Immortalized control, FSHD1, FSHD2 and control-derived mutant skeletal myoblast cells were grown in high glucose DMEM (Gibco) supplemented with 20% FBS (Omega Scientific, Inc.), 1% Pen-Strep (Gibco), and 2% Ultrasor G (Crescent Chemical Co.). Immortalization and single cell clone isolation of primary FSHD1 myoblasts were performed as previously described for Control and FSHD2 myoblasts (60). Upon reaching 80% confluence, myoblast differentiation was induced by using high glucose DMEM medium supplemented with 2% FBS and ITS supplement (insulin 0.1%, 0.000067% sodium selenite, 0.055% transferrin; Invitrogen). Fresh differentiation media was changed every day.

### Antibodies

Immunofluorescence (IF) or Western blot (WB) is performed using antibodies specific for SMCHD1 (NBP1-49968, Novus Bio.), DUX4 (NBP2-12886, Novus Bio.), LEUTX (PA5-59595, Thermofisher), Actin (A4700, Sigma), Histone H3 (ab18521, Abcam), and Histone H3.X/Y (MABE243I, Sigma).

### Immunofluorescent staining

Staining was performed as previously described (61). Briefly, cells grown on coverslips were fixed in 4% paraformaldehyde for 10 min at room temperature, permeabilized with 0.5% Triton X-100 in PBS, and blocked in blocking buffer (0.02% saponin, 0.05% NaN_3_, 1% BSA, 4% horse serum and 0.1% gelatin in PBS) for 15 min at 37°C. The coverslips were incubated overnight with primary antibodies at 4 °C followed by three PBS washes, then incubated with fluorescent secondary antibody for 30 min at 37°C, washed with PBS 3 times, counter-stained with DAPI, and mounted with Prolong Diamond Antifade Mountant. Images were acquired with a Zeiss LSM510 confocal laser microscope.

### Western blotting

Cells were lysed in 2X Laemlli Buffer with 4% beta-mercaptoethanol, sonicated, boiled, and separated by 4%–20% TEO-Tricine gel (Abcam). Then the samples were transferred to nitrocellulose membranes, blocked with Pierce Protein-Free T20 (PBS) Blocking Buffer (Thermo Fisher Scientific), and blotted with the desired antibodies. Horseradish peroxidase-conjugated anti-mouse-IgG (Promega), anti-rabbit-IgG (Promega), or anti-rat-IgG (Abcam) were used as secondary antibodies. Immunoblots were developed with SuperSignal West Pico Chemiluminescent Substrate (Thermo Fisher Scientific). Images were acquired using the Image Analyzer (LAS-4000, Fujifilm).

### RNA isolation and quantitative real-time RT-PCR (RT-qPCR)

RNA was extracted using RNeasy Plus Mini kit (Qiagen, Cat No. 74134), and complementary DNA (cDNA) was made using 500ng of total RNA with SuperScript IV VILO Master (Thermo Fisher Scientific, Cat No. 11756050) following the manufacturer’s instructions. qPCR was performed by using AzuraView GreenFast qPCR Blue Mix LR (Azura Genomics Inc., Cat No. AZ-2320). The genes and their corresponding PCR primers are listed in Supplemental Table S4. For some genes, the expression was detected by probe qPCR, which was performed by using TaqMan Fast Advanced Master Mix (Thermo Fisher Cat No. 4444557). The commercially available TaqMan Gene Expression Assay probes (Thermo Fisher Scientific) were listed in Supplemental Table S5.

### RNAScope in situ hybridization

RNAscope was performed using RNAscope Multiplex Fluorescent Reagent Kit v2 (Advanced Cell Diagnostics, Inc., Cat. No. 323100) according to the manufacturer’s protocol as previously described (37). The following RNAScope probes (Advanced Cell Diagnostics, Inc.) were used: *LEUTX* probe set (Hs-LEUTX-C2, Cat. No. 547251-C2), 6ZZ *DUX4fl* probe set (HS-DUX4-O6-C1, Cat. No. 546151), 4ZZ *DUX4fl* probe set (Hs-DUX4-O7-C2, Cat. No. 1089191-C2), and 2ZZ *DUX4fl* probe set (Hs-DUX4-O8-C3, Cat. No. 1089201-C3). The 4ZZ and 2ZZ *DUX4fl* probes target 701-1388 and1481-1697 of *DUX4fl* mRNA (NM_001306068.2) respectively (Figure 5E).

### 5-Azacytidine (5AzaC) and 5-Aza-2’-deoxycytidine (5AzadC) treatment

Six μM of 5AzaC (Sigma-Aldrich, A2385) was added to myoblasts at ~70% confluency for 48h. With or without differentiation, the cells were harvested or fixed at indicated day after the drug treatment for subsequent RNA-seq, ChIP-qPCR, RT-qPCR, MeDIP and IFA analyses. Alternatively, myoblasts were incubated with 6 μM of 5AzadC (Sigma, A3656) for 24h, followed by 2 days release. To assess the SMCHD1 depletion effect, cells were treated with 5 μM of 5AzaC for 48 hours. For RNA-seq experiment, myoblasts were allowed to grow for additional 48 hours before collection or before changing to differentiating media. For SMCHD1 mutant maintenance experiment, cells were allowed to recover for 48 hours before collection.

### ShRNA depletion or overexpression of proteins using lentiviral systems

The shRNA plasmids for SMCHD1 (5’-TTATTCGAGTGCAACTAATTT-3’, TRCN0000253777), DUX4 (5’-AGATTTGGTTTCAGAATGAGA-3’, TRCN0000421072) and a control shRNA (shCTRL, 5’-CAACAAGATGAAGAGCACCAA-3’, SHC002), were obtained from the Sigma Mission library. The shRNA pLVshH3.X/Y (5’-GCGGGAAATCAGAAAGTAC-3’, the siH3.X/Y targeting sequence in previous paper (47)), pLVshDUX4 (5’-GGCAAACCTGGATTAGAGTT-3’ (18)), and corresponding pLVshControl (5’-CCTAAGGTTAAGTCGCCCTCG-3’) were synthesized and cloned in pLV(shRNA)-Puro-U6 vector by VectorBuilder. To construct the H3.X and LEUTX overexpression lentiviral plasmids pLVX_H3.X and pLVX_LEUTX, human H3.X and LEUTX ORF sequences were amplified from FSHD1 myotube cDNA using primer pairs H3.X_PCR and LEUTX_PCR respectively, and cloned into pLVX Lentiviral vector (Addgene, plasmid #135182). An empty pLVX vector was used as a negative control. Lentivirus packaging, transduction and puromycin selection were performed as previously described with slight modifications (37). Briefly myoblasts were infected twice at 48 hour and 24 hours prior to differentiation. The differentiated cells were harvested at days 3-7 when 60-80% cells fused to form myotubes.

### Detection the ratio of 4qA- and 10q-specific nucleotide polymorphisms (SNPs) using Amplicon sequencing

Input and MeDIP DNA were amplified by PCR using Phusion DNA Polymerase in two steps: The first PCR primers (D4Z4_seqPCR) are derived from the 4q/10q specific Q-PCR primers (24) with adapters for the 2nd PCR primers attachment. The 2nd PCR and Amplicon sequencing were performed following the previous paper (62) with modifications. The second PCR primer pair (Illumina_seqPCR) was used to attach Illumina adaptors and to barcode samples. Amplification was carried out with 18 cycles for the first PCR and 24 cycles for the second PCR. Resulting amplicons from the second PCR were gel extracted, quantified, mixed and sequenced using NovaSeq6000 (Illumina). The ratio of 4qA/B - and 10q-derived D4Z4 sequences was calculated based on the reads number of their specific SNPs.

### Pulsed-field gel electrophoresis (PFGE) and Southern blotting for 4q/10q D4Z4 repeat array length analysis

PFGE and southern blotting were performed as described (59) with slight modifications. Briefly, suspended myoblasts were mixed with melted 1% UltraPure Low Melting Point Agarose (Bio-Rad) at 37°C to form plugs, each containing ~1.5 x10^6^ cells. The plugs were digested with pronase (Sigma), rinsed, and treated with enzymes (EcoRI plus HindIII or EcoRI plus BlnI, Roche). The digested DNA in plugs was subjected to PFGE.

Electrophoresis was done in CHEF-DR III system (Bio-Rad) at 6 V/cm and 15°C for 13 h, with the switch time increasing linearly from 1 to 6 sec. The gel was washed in 0.25 M HCl for 30 minutes to depurinate DNA fragments, rinsed with H_2_O, washed in denaturation buffer (0.6 M NaOH, 0.4 M NaCl) for 30 minutes, and transferred in that solution to a Biodyne B Membrane (KPL). The membrane was neutralized, cross-linked by UV irradiation, and was probed with the 1-kb 4q/10q specific probe described in the paper (59). Southern blots were visualized using Typhoon scanner (GE Healthcare).

### ChIP-qPCR Analysis

Around 3 x 10^6^ cells were crosslinked with 1% formaldehyde for 10 minutes and quenched with 0.125 M glycine for 5 minutes. Cells were washed with ice-cold PBS and harvested in cell lysis buffer (5 mM PIPES pH 8.0, 85 mM KCl, 0.5% NP40, protease inhibitors). Nuclear extract was collected in RIPA buffer (1 % NP-40, 0.5% Sodium Deoxycholate, 0.1% SDS, protease inhibitors). Nuclei were sonicated for 20 cycles (30 seconds on and off) using the Bioruptor (Diagnode) to obtain fragment length around 150-500 bp. Chromatin extract was quickly spun down to remove cell debris. Around 20 μg or 10 μg of chromatin were incubated overnight at 4°C with 10 μg H3.X/Y or 2.5 μg H3K9me3 antibody respectively (Active Motif-61161, Abcam-ab8898). Chromatin-antibody extracted with incubated with protein G Dynal beads (Thermo Fisher Scientific, 10003D) for an hour. The mixture was washed with Lithium Chloride 5 times and TE buffer once on ice. Chromatin was eluted with 0.1 M NaHCO3 and 1% SDS. Proteinase K was added to chromatin and the mixture was proceeded to reverse-crosslinking for 2 hours at 55°C. DNA was purified with QIAquick PCR Purification Kit (Qiagen). ChIP DNA was quantified by qPCR with specific primers.

### MeDIP

Cells were washed with PBS and harvested in SDS lysis buffer (1% SDS, 10 mM EDTA, 50 mM Tris HCl, pH 8.1). Nuclear extract was sonicated with 3 cycles (30 seconds on and off). Chromatin extract was quickly spun down and supernatant was collected. Chromatin was added with Proteinase K and incubated for 2 hours at 55°C. DNA was purified with QIAquick PCR Purification Kit (Qiagen). Between 0.5-1 μg of chromatin was proceeded to MeDIP (Methylated DNA Immunoprecipitation) according to the manufacture protocol (EpiMark® Methylated DNA Enrichment Kit, New England Biolabs). MeDIP DNA was quantified by qPCR with specific primers and sequencing.

### RNA-seq and data processing

Total RNA was extracted by using the RNeasy kit (QIAGEN). Between 12-50ng of RNA was converted to cDNA using the Smart-Seq 2 protocol (38). DNA libraries were constructed using Nextera DNA Flex Library Prep Kit (Illumina). DNA samples were sequenced on the Illumina NextSeq500 platform using paired-end 43 bp mode with around 15 million reads per sample. RNA-seq raw reads were aligned with STAR (version 2.5.1b) using human genome reference hg38. Alignment default parameters were applied except with a maximum of 10 mismatches per pair, a ratio of mismatches to read length of 0.07, and a maximum of 10 multiple alignments. Read count was performed using RSEM (version 1.3) by defaults with gene annotations from GENCODE v28, and raw read counts were extracted for downstream analysis. Genes were filtered based on raw counts with at least 2 counts in at least 2 samples. Raw count was normalized by TMM in EdgeR and then converted to TPM (transcript per millions). Differential genes were calculated using the cut off *P-*values =< 0.01 and Log_2_ Fold Change >=1.5.

### Statistical Analyses

Microsoft Excel software was used to perform statistical analyses on data from three independent experiments. Statistical comparisons were made using the unpaired Student’s t-test and Wilcoxon’s t-test. Statistical significance between two samples was determined by a p value of less than 0.05. Error bars shown mean ± SD.

## Supporting information

A list of DEGs in all three mutant types

A list of ranked high genes (500) in PC1 and PC2 in Figure 2A

A list of genes in different GO terms in Figure 3A and B

The sequences of the primers used in this study

TaqMan Gene Expression Assay probes

## Acknowledgments

pH082 pU6-gRNA2.0-GFP is a kind gift from Dr. Gerd A. Blobel, Children’s Hospital of Philadelphia. The authors wish to acknowledge the support of the Chao Family Comprehensive Cancer Center Optical Biology Core (LAMMP/OBC) Shared Resource.

## Data and materials availability

RNA-seq data were deposited to dbGAP (phs002554.v2.p1). All other data are available in the main text or the supplementary materials.

---

**Fig. S1.**
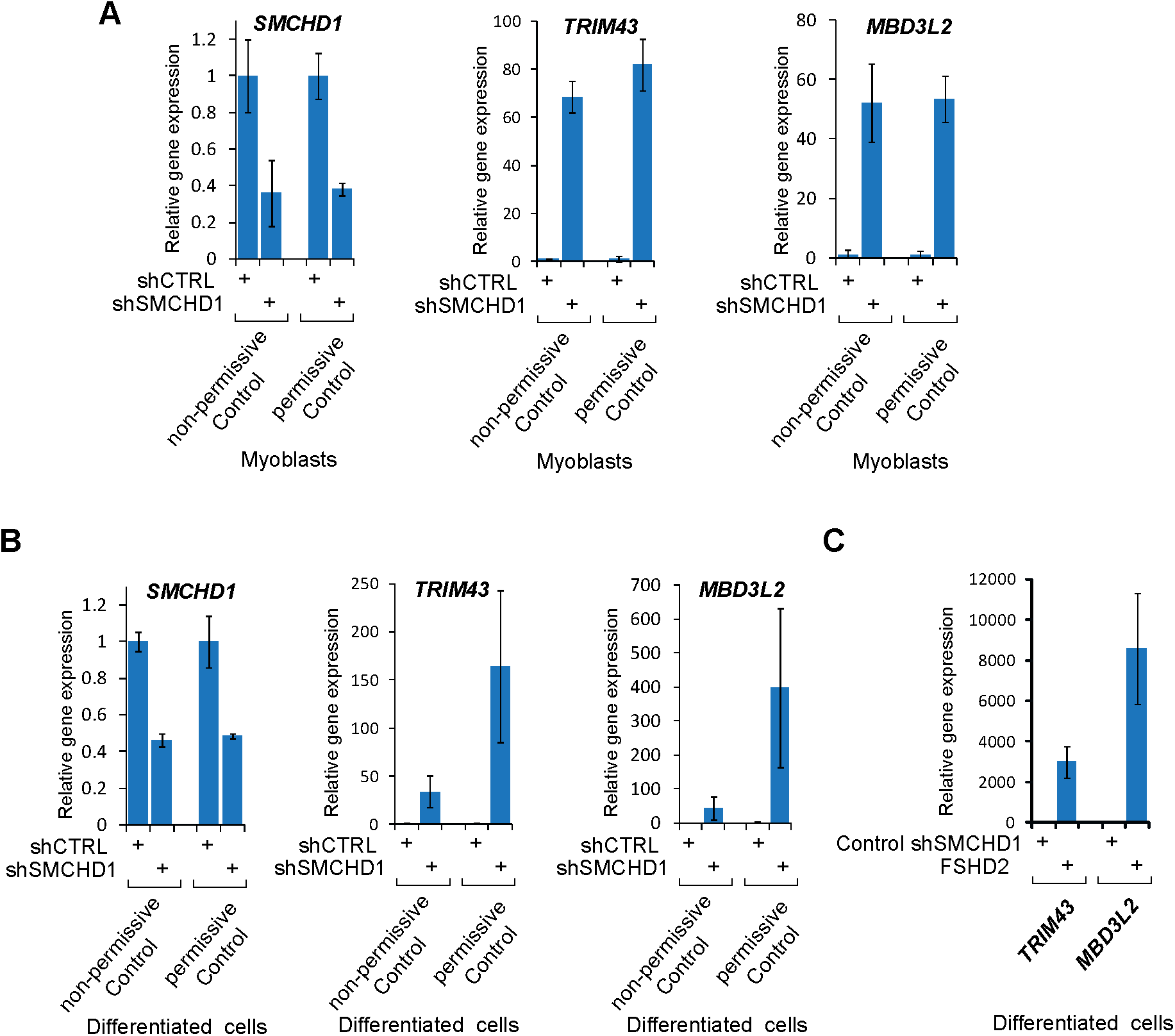
Comparison of transient depletion of SMCHD1 in myoblasts with permissive or non-permissive haplotype. **A.** Upregulation of DUX4 target genes are comparable in both haplotypes at the myoblast stage. RT-qPCR analyses of SMCHD1 depletion efficiency and expression of two DUX4 target genes (TRIM43 and MBD3L2) in permissive or non-permissive myoblasts treated with control shRNA and shRNA specific to SMCHD1. Data are presented as fold change in expression relative to the respective control shRNA-treated myoblasts. **B.** Similar experiments as in (A) in myotubes. Following lentiviral shRNA infection, myoblasts were differentiated into myotubes for 3 days and RT-qPCR analyses were performed for *SMCHD1, TRIM43* and *MBD3L2* as indicated. **C.** RT-qPCR comparison of shSMCHD1-treated myotubes and FSHD2 patient myotubes on day 3 of differentiation, demonstrating the vast overexpression of TRIM43 and MBD3L2 in patient myotubes.

**Fig. S2.**
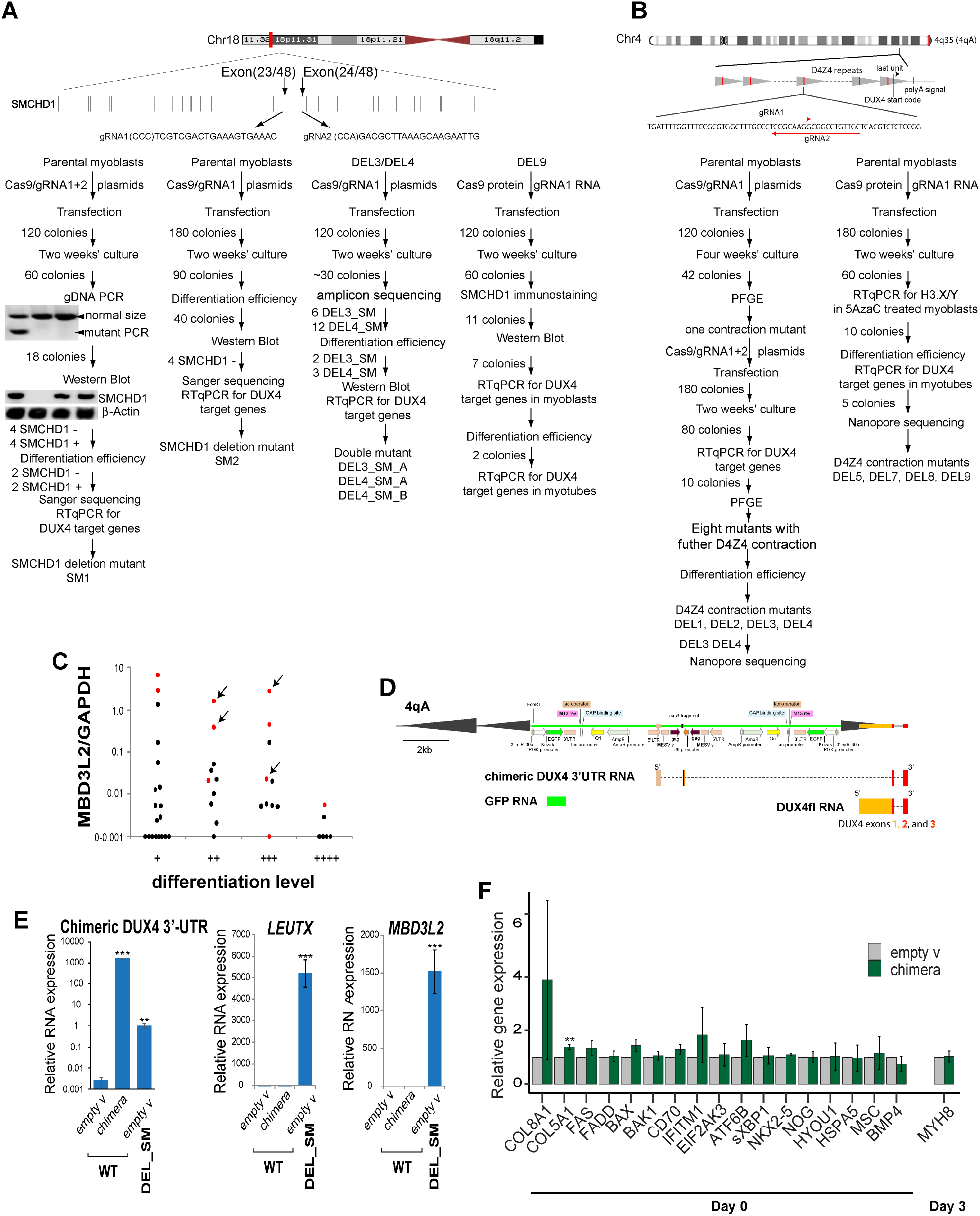
Applications of CRISPR/Cas9 to generate FSHD modeling cells. **A.** A scheme of the experimental procedure for generating and isolating single clones for *SMCHD1* knockout (SM) mutants. gRNA targeting sequences and PAM sequences(red) are shown. Two sets of screening for SM mutants and two sets of screening for double (DEL_SM) mutants were performed. *SMCHD1* mutants were confirmed by western blot and gRNA targeting region genomic DNA sequencing. **B.** A scheme of the experimental procedure for screening and isolating single clones for D4Z4 deletion (DEL) mutants. For the D4Z4 repeat unit number and haplotype, the sizes of markers and their migration distance in the PFGE can be used to determine the size of the D4Z4 fragments using a standard curve Excel. Nanopore sequencing was also used to sequence chromosome 4 (chr4) D4Z4 region and chromosome 10 (chr10) D4Z4 region. **C.** DUX4 target gene expression level and differentiation efficiency of potential single contraction mutant colonies. The cell lines marked with red dot were subjected to PFGE. Arrows indicate the cell lines used for other experiments in this paper. All the cell lines, which have the best differentiation efficiency, showed low MBD3L2 qPCR signal. One of them was confirmed no further contraction by using PFGE. All the tested cell lines with higher MBD3L2 signal, were deletion mutants. **D**. Detailed schematic diagram of the Cas9/gRNA plasmids insertion in 4qA allele, the adjacent D4Z4 units, and the RNAs transcribed from this region. **E.** Left: artificial overexpression of a chimeric DUX4 3’UTR RNA in parental WT cells and detection of the endogenous transcript in DEL_SM mutant cells by RT-qPCR. Middle and right: overexpression of chimeric DUX4 3’UTR induces no *LEUTX* or *MBD3L2* expression in WT cells. Expression of these genes in DEL_SM cells transfected with an empty vector is shown for comparison. **F.** Similar overexpression of chimeric DUX4 3’UTR in myoblasts and myotubes had no effect on the genes found to be altered in mutant cells as in Figure 3C.

**Fig. S3.**
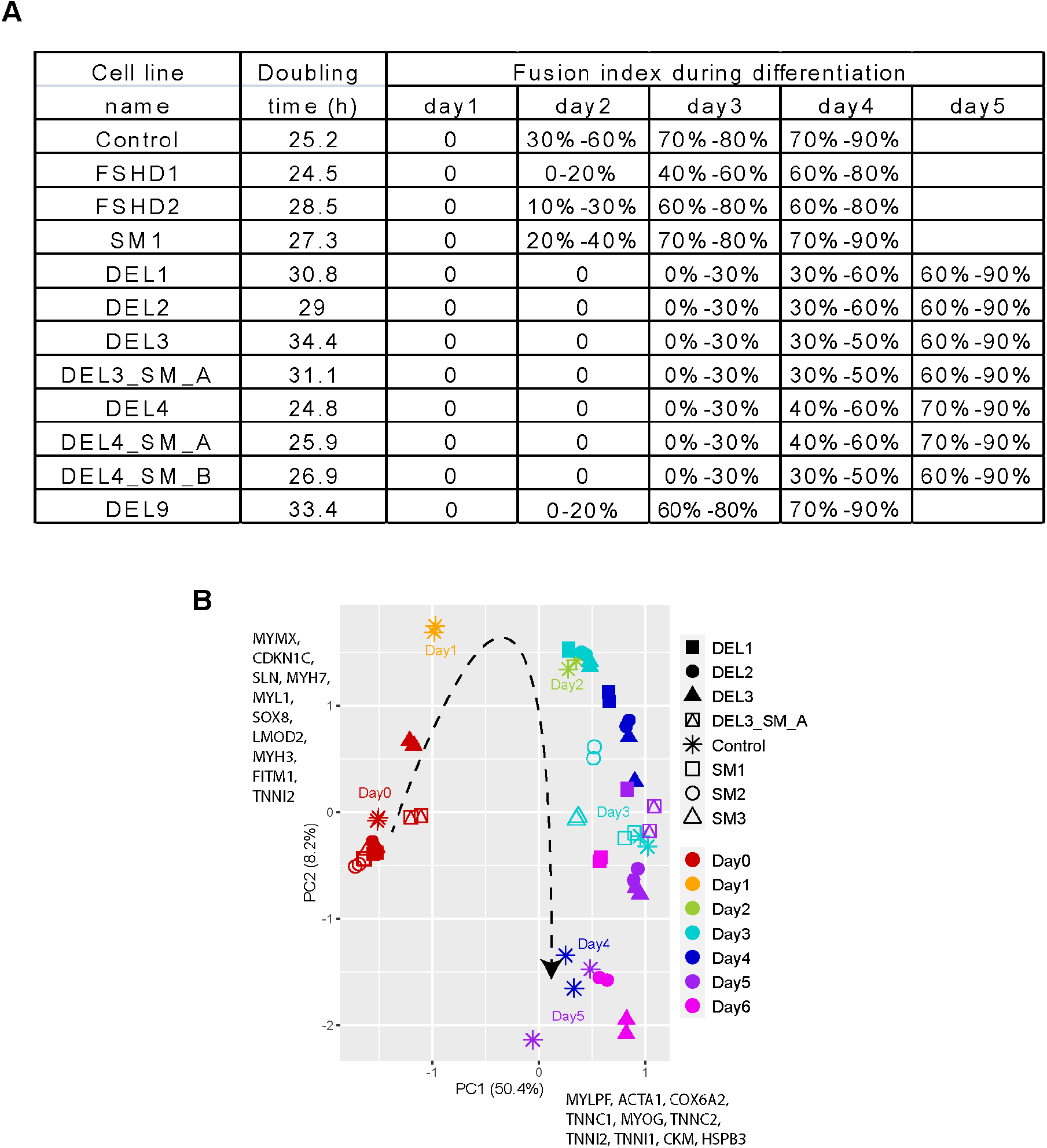
Assessment of proliferation and differentiation of mutant myocytes. **A.** A table summary for doubling time and differentiation efficiency. Total myoblast cell number in a field was counted when the cells were confluent one day before fusion. After fusion, the mononuclear cells number in this filed was counted at different day. Differentiation efficiency was then determined by [the initial cell number] – [the number of mononucleated cells left on each day] / [the initial cell number]. **B.** Principal component analysis of control and FSHD mutants along differentiation days from RNA-sequence experiment. The arrow indicates the trajectory of the control differentiation time course from Day 0 to Day 5. Both PCA1 and PCA2 explain variance across differentiation. Colors indicate days of differentiation and shapes indicate cell lines. The top 10 genes are shown for each component.

**Fig. S4.**
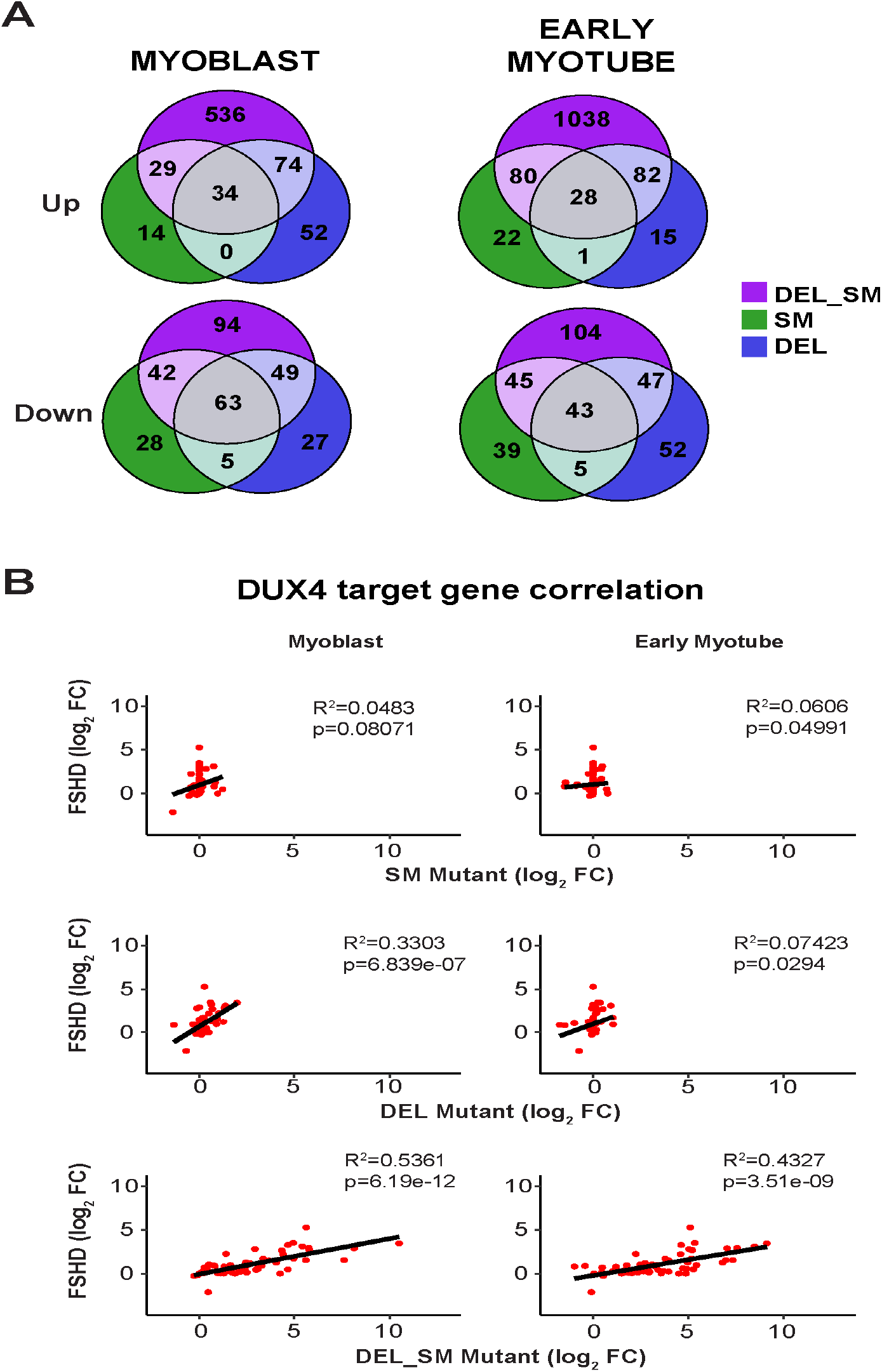
RNA-seq data analyses of three types of mutants and patient cells. **A.** Venn diagrams show the intersection of DEGs compared to control for each mutant type (SM, DEL and DEL_SM mutants) at myoblast and early myotube. **B.** R square correlation plots for DUX4 targets (64 total) between both FSHD patient lines and each mutant type (SM, DEL, DEL_SM mutants) at myoblast and early myotube. *P*-values indicate significant correlations.

**Fig. S5.**
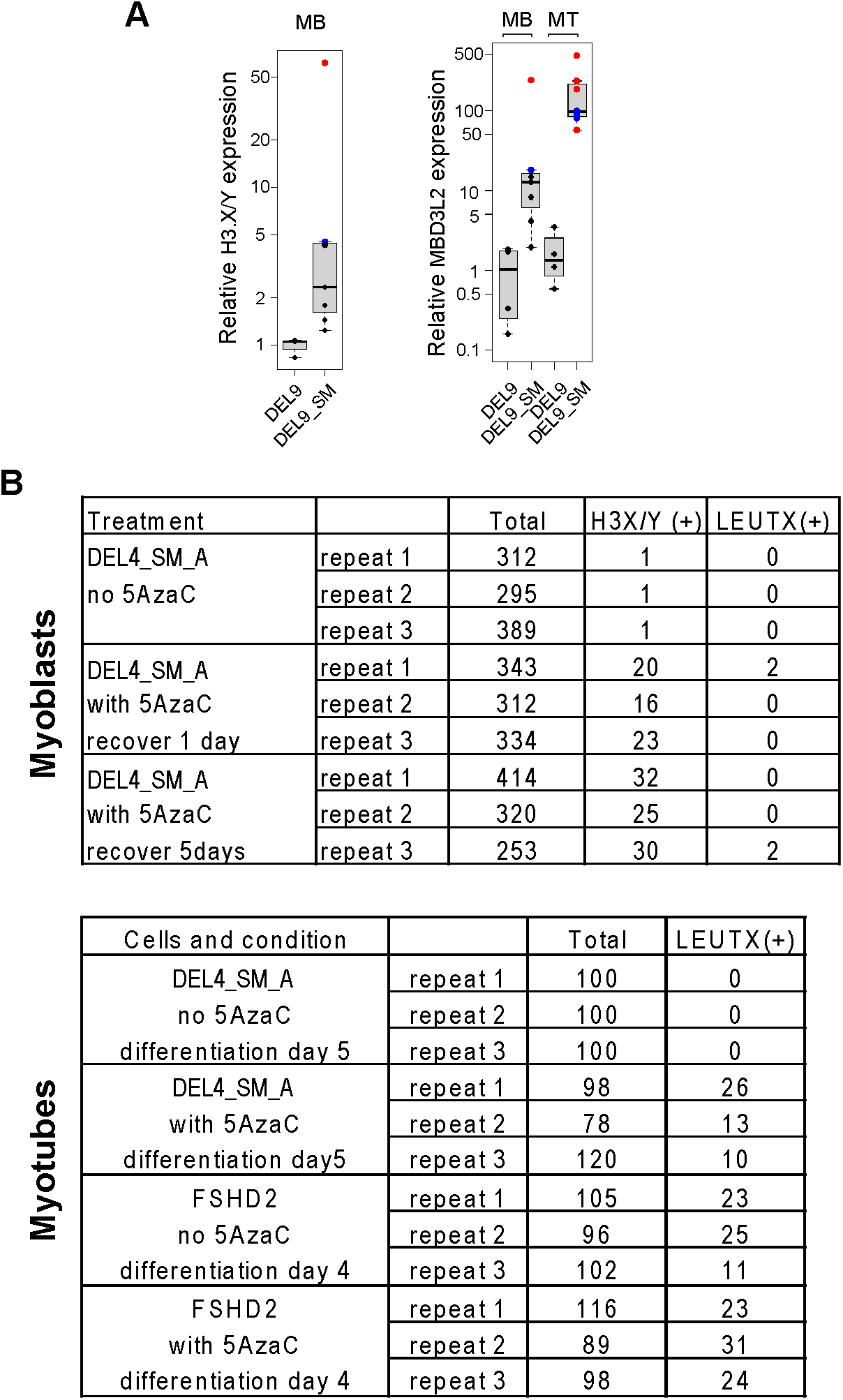
Synergistic effect of double mutations and the effect of 5AzaC treatment. **A.** The effect of SMCHD1 mutation on H3.X/Y and MBD3L2 expression in DEL9 mutant myoblasts (MB) and myotubes (MT) as indicated. Two double mutants (red and blue) with similar differentiation efficiency as DEL9, were used for the MT data. **B.** The number of myotubes expressing H3.X/Y or LEUTX proteins without or with 5AzaC treated myoblasts or myotubes as indicated. Total number of cells counted as well as positive cells in triplicate experiments are shown.

**Fig. S6.**
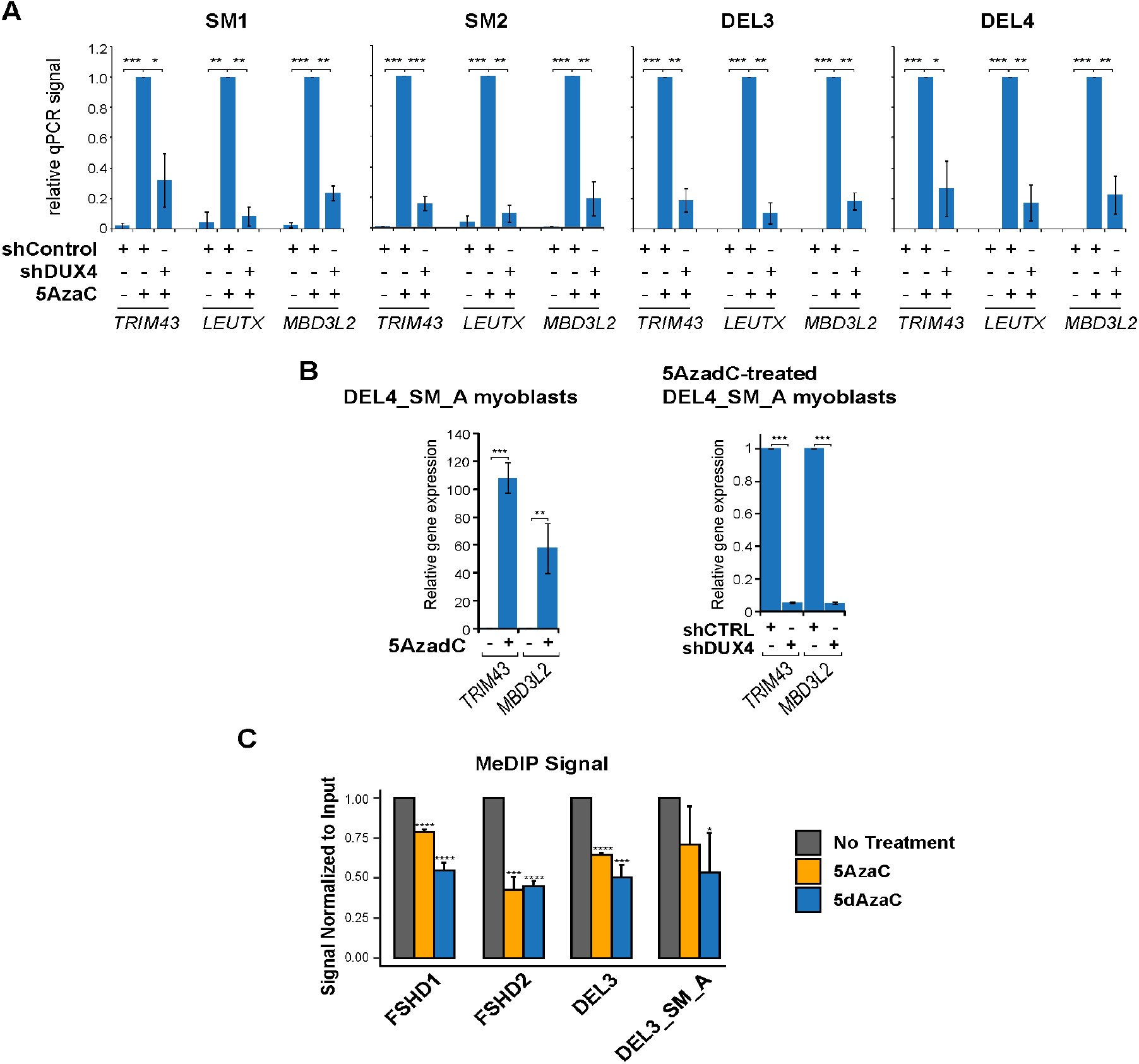
Analysis of 5AzaC and 5AzadC treatment. **A.** 5AzaC induced upregulation of DUX4 target genes in mutant cells was DUX4-dependent. Cells were treated with 5AzaC, infected with lentivirus containing shCTRL or shDUX4, and induced differentiation same as Figure 6D. For each cell line, DUX4 target genes expression level after DUX4 depletion was shown as fold difference compared to the control. Data are presented as mean ± SD; **p<0.01, by one-tailed student’s t-test. **B.** DUX4 target genes were upregulated by 5AzadC treatment in double mutant myoblasts (left panel). This upregulation was inhibited by DUX4 depletion (right panel). Cells were treated with 5AzadC for 24 h and harvested for gene expression analysis 2 days after release from 5AzadC. For the DUX4 depletion experiment, the cells were infected with shCTRL or shDUX4 lentivirus one day before 5AzadC treatment. **C.** The effect of 5AzaC and 5dAzaC treatment on FSHD, single deletion and double mutant at Day 5 myotubes. Cells were treated with 5AzaC or 5dAzaC for 48 hours before differentiation. MeDIP analysis was performed to confirm the inhibition of DNA methylation at D4Z4. Significant inhibitions are indicated by the asterisks.

**Fig. S7.**
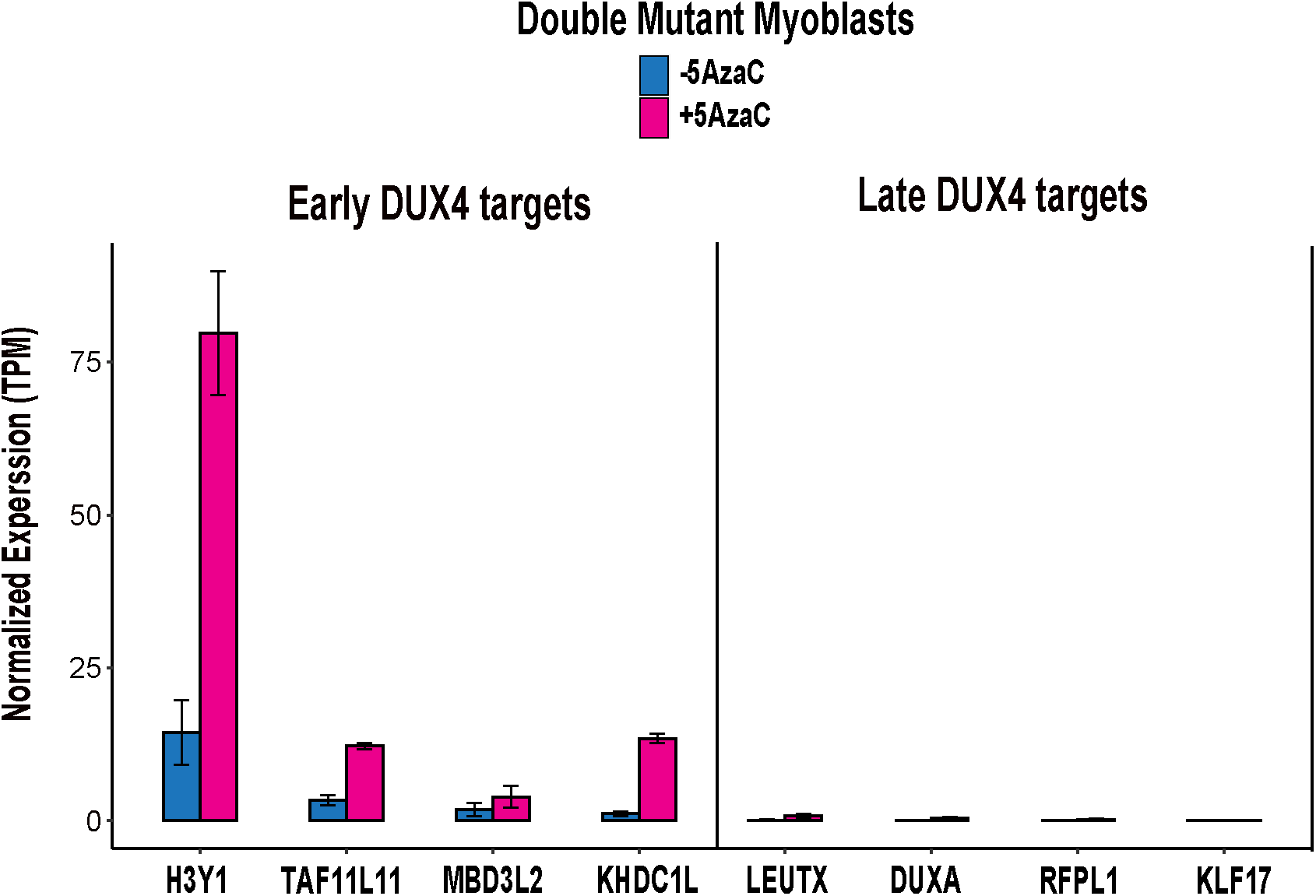
Comparison of early and late DUX4 target genes in double mutant myoblasts with or without 5AzaC treatment. In contrast to the early target genes, the late target genes are refractory to 5AzaC treatment at the myoblast stage.

